# Major revisions in pancrustacean phylogeny with recommendations for resolving challenging nodes

**DOI:** 10.1101/2022.11.17.514186

**Authors:** James P. Bernot, Christopher L. Owen, Joanna M. Wolfe, Kenneth Meland, Jørgen Olesen, Keith A. Crandall

## Abstract

The clade Pancrustacea, comprising crustaceans and hexapods, is the most diverse group of animals on earth, containing over 80% of animal species. It has been the subject of several recent phylogenomic analyses, but despite analyzing hundreds of genes, relationships within Pancrustacea show a notable lack of stability. Here, the phylogeny is estimated with expanded taxon sampling, particularly of malacostracans, using a precise tree-based approach to infer orthology. Our results show that small changes in taxon sampling have a large impact on phylogenetic estimation. By analyzing only shared orthologs between two slightly different taxon sets, we show that the differences in the resulting topologies are due to the effects of taxon sampling on the phylogenetic reconstruction method, not on ortholog identification. We compare trees resulting from our phylogenomic analyses with those from the literature to explore the large tree space of pancrustacean phylogenetic hypotheses and find that statistical topology tests reject the previously published trees in favor of the ML trees produced here. Our results reject several clades including Caridoida, Eucarida, Multicrustacea, Vericrustacea, and Syncarida. We recover a novel relationship between decapods, euphausiids, and syncarids that we refer to as the Syneucarida. With denser taxon sampling, we find Stomatopoda sister to this clade, which we name Stomatocaridea, dividing Malacostraca into three clades: Leptostraca, Peracarida, and Stomatocaridea. A new Bayesian divergence time estimation is conducted using 13 vetted fossils. We review our results in the context of other pancrustacean phylogenetic hypotheses and highlight the key taxa to sample in future studies.

## Introduction

The clade Pancrustacea (Crustacea + Hexapoda) is arguably the most successful group of animals on earth. It comprises 1,236,586 species, contains more than 80% of extant animal diversity (Roskov et al. 2020), and includes nearly half of all animal biomass on the planet (Bar-On et al. 2018). Pancrustaceans have been a dominant component of earth’s ecosystems for nearly 600 million years (Wolfe et al. 2016). The group includes 55 orders of crustaceans and 31 orders of hexapods (Bracken-Grissom and Wolfe 2020). Many of the most economically important species on earth are pancrustaceans including bees, mosquitos, krill, and numerous other taxa with key positions in terrestrial and aquatic food webs. The morphological diversity of body plans in this group is unparalleled among animals (Fig. 1); for example, ranging from minute 70 µm tantulocarid larvae (Huys et al. 1993; Petrunina et al. 2018) to Japanese spider crabs with a leg span up to 3.7 meters (McClain et al. 2015). While the species diversity of Pancrustacea is dominated by hexapods, the ~64,000 species of non-insect crustaceans (WoRMS 2022) make up most of the phylogenetic diversity and morphological disparity of Pancrustacea. The non-hexapod Pancrustacea diversity is composed of ten classes of crustaceans: Branchiopoda, Cephalocarida, Copepoda, Ichthyostraca (i.e., Branchiura + Pentastomida), Malacostraca, Mystacocarida, Ostracoda, Remipedia, Tantulocarida, and Thecostraca (WoRMS 2022). Despite a number of recent pancrustacean phylogenomic studies (Regier et al. 2008; Regier et al. 2010; Andrew 2011; von Reumont et al. 2012; Oakley et al. 2013; Rota-Stabelli, Lartillot, et al. 2013; Schwentner et al. 2017; Schwentner et al. 2018; Lozano-Fernandez et al. 2019), the relationships among classes have been particularly challenging to reconstruct for a number of reasons. First, the most recent common ancestor of Pancrustacea has been estimated to be greater than 500 million years ago (Ma) (Oakley et al. 2013; Rota-Stabelli, Daley, et al. 2013; Schwentner et al. 2017; Wolfe 2017), prior to the occurrence of recognizable arthropod fossils (Wolfe et al. 2016; Daley et al. 2018). The difficulty with confidently estimating deep-time relationships among taxa of this age is further compounded by the suggestion that these relationships are part of a rapid radiation. Evolutionary relationships that are part of an ancient rapid radiation are some of the most difficult to resolve due to their short internal branch lengths and age (Fishbein et al. 2001; Rokas and Carroll 2006; Whitfield and Lockhart 2007; One Thousand Plant Transcriptomes Initiative 2019). Generally, to obtain robust nodal supports for rapid radiations, more genes are sequenced to add as much information to those short branches as possible, but this is difficult for many pancrustaceans; many lineages are rarely collected, small bodied, and often have large genomes (Alfsnes et al. 2017; Bracken-Grissom and Wolfe 2020), so maximizing sequence data through whole genome sequencing remains challenging and costly. In addition, pancrustaceans are known to have short exons (e.g., Owen et al. 2020), which yield shorter alignments with fewer parsimony informative sites to resolve branches in phylotranscriptomic studies. Furthermore, short exons are vulnerable to weak signal:noise ratios, which can be particularly problematic for multispecies coalescent models (Huang et al. 2010; Bayzid and Warnow 2013; Patel et al. 2013; DeGiorgio and Degnan 2014; Mirarab et al. 2014; Lanier and Knowles 2015; Mirarab and Warnow 2015; Xi et al. 2015). Unfortunately, a robust and comprehensive morphology-based phylogeny is also not available for Pancrustacea due to challenges with ancient, rapid radiations resulting in extreme morphological variation between groups and lacking or ambiguous support for many clades (Wolfe 2017; Bracken-Grissom and Wolfe 2020).

**Figure 1.**
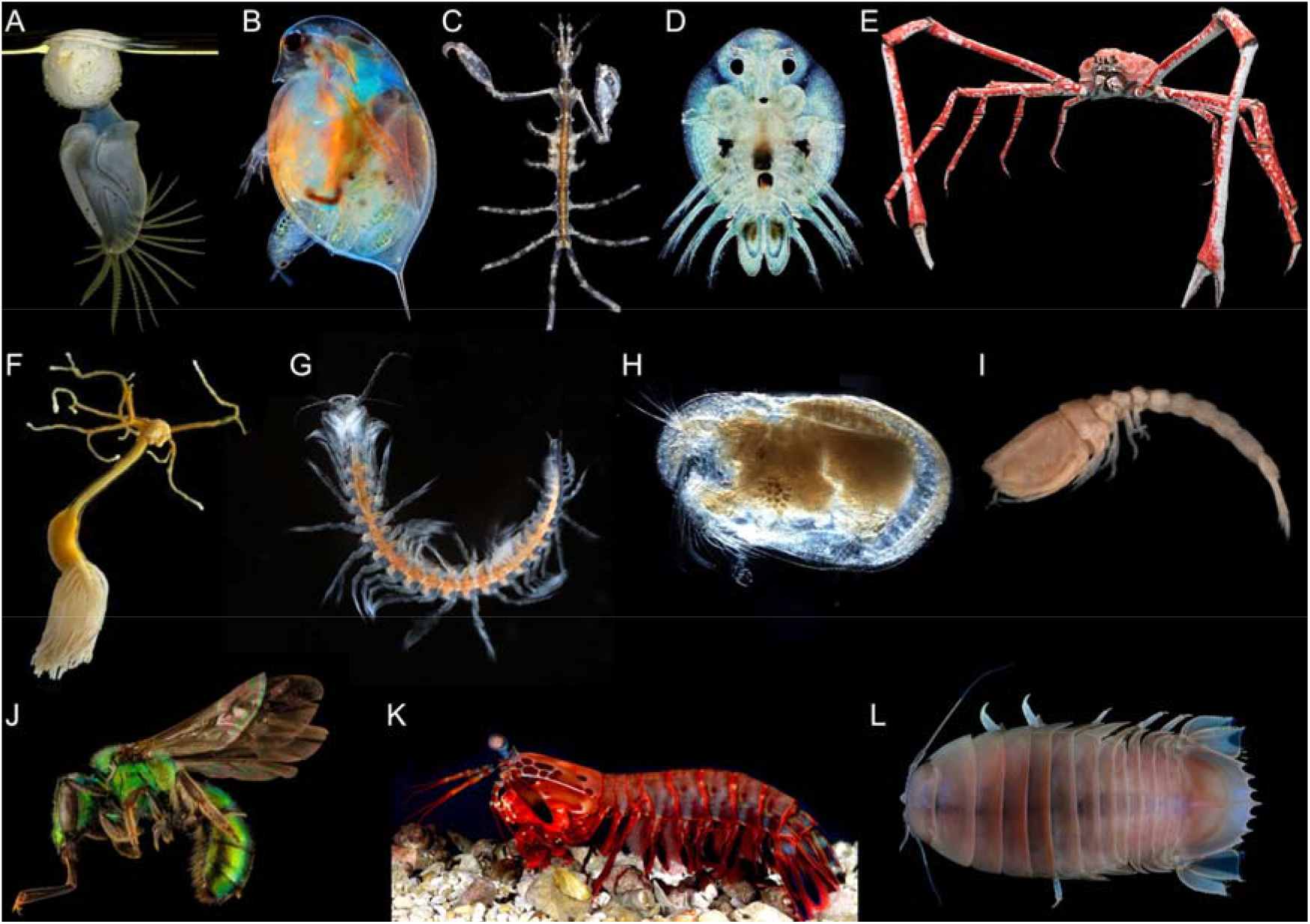
Morphological diversity of select pancrustaceans. (A) Buoy barnacle (Cirripedia). (B) *Daphnia* sp. (Branchiopoda). (C) Skeleton shrimp (Amphipoda). (D) *Argulus* sp. (Branchiura). (E) Japanese spider crab (Decapoda). (F) Parasitic copepod (Copepoda). (G) *Lasionectes entrichoma* (Remipedia). (H) Seed shrimp (Ostracoda). (I) Comma shrimp (Cumacea). (J) Sweat Bee (Hexapoda). (K) Mantis shrimp (Stomatopoda) (L) Giant isopod (Isopoda). Image credits: (A) David Fenwick, (B) Marek Miś (C) David Fenwick, (D) Andrei Savitsky, (E) Michael Wolfe and Hans Hillewaert, (F) Geoff Boxshall, (G) Jørgen Olesen, (H) Anna Syme, (I) Hans Hillewaert, (J) USGS Bee Inventory and Monitoring Lab, (K) Roy L. Caldwell, (L) Chan T. Y. & Lin C.W.

Historically, a variety of analytical approaches have been employed to overcome the difficulties in estimating a robust Pancrustacea phylogeny. These include a variety of phylogenetic methods (e.g., Dayhoff matrix recoding, partitioned analyses vs site-heterogeneous models) as well as alternative methods of ortholog selection (e.g., sequence similarity vs tree based). Despite the many different phylogenomic analyses, few have focused on taxon sampling; instead, a small number of novel taxa are typically added for each study while most data are re-used from public databases. Currently, nearly half of the 55 crustacean orders, many of which are rare taxa, have never been included in a multi-gene phylogeny, and relationships of the orders that have been sampled have often been unstable; phylogenetic relationships frequently change when sequence data and taxa change (Mallatt and Giribet 2006; Regier et al. 2008; Regier et al. 2010; Andrew 2011; von Reumont et al. 2012; Oakley et al. 2013; Rota-Stabelli, Lartillot, et al. 2013; Schwentner et al. 2017; Schwentner et al. 2018; Lozano-Fernandez et al. 2019).

Despite being integral in systematic study design, taxon sampling has received less focus relative to phylogenomic methodology in studies of pancrustacean phylogeny. Prior to the phylogenomics era, there was extensive debate over whether it is more important to sample more taxa or more nucleotide sites (reviewed in Nabhan and Sarkar 2012). In general, those arguing in favor of taxon sampling cite studies that have demonstrated that well-sampled phylogenies lead to more accurate topologies (e.g., Hedtke et al. 2006; Heath et al. 2008), more accurate branch lengths (e.g., Hugall and Lee 2007), and reduce the number of long branches that may contribute to long branch attraction (e.g., Hendy and Penny 1989; Poe 2003). To date, no phylogenomic study has demonstrated the effects of taxon sampling in relation to the Pancrustacea. This is important because the lack of comprehensive taxonomic coverage at the ordinal rank (described above) may impact topological accuracy and studies have shown that taxon sampling does impact tree topology even when thousands of loci are used. For example, Betancur-R et al. (2019) and Branstetter et al. (2017) demonstrated that taxon sampling and density both contribute to accuracy when using genomic data. Despite its significance in molecular systematics, taxon sampling has received relatively little attention in the phylogenomics era and its effects have not been tested with respect to the Pancrustacea phylogeny.

In addition to these challenges, many questions remain unanswered regarding the evolutionary relationships within Pancrustacea. Key areas of contention include: the position of Hexapoda, Ostracoda monophyly, interrelationships of Multicrustacea (Copepoda, Malacostraca, Thecostraca), and relationships within the most speciose crustacean class, Malacostraca. The evolutionary relationships of most non-hexapod pancrustacean orders also remain unresolved and many orders have yet to be sampled with genome scale data. To address these questions, we estimate pancrustacean relationships with increased taxon sampling using 106 transcriptomes and 14 genomes (Table S1), and a tree-based approach for ortholog selection, which has been shown to improve phylogenetic reconstruction (Dunn et al. 2013; Yang and Smith 2014; Ballesteros and Hormiga 2016; Smith and Pease 2017). With the resulting phylogeny, we examine deep level relationships of pancrustaceans and estimate the timing of their divergence using 13 fossils as calibration points. We also compare these results to two different taxon sampling schemes from prior versions of this study; these results demonstrate that, even in the context of hundreds of orthologs, taxon sampling and outgroup selection have major impacts on the phylogenetic tree search under all methods used here. By controlling for ortholog selection, we show that the differences in topology among our most similar taxon sets are driven by the effects of taxon sampling on the phylogenetic tree search rather than on the ortholog identification process. We review our results in the context of all other pancrustacean phylogenomic studies to identify areas of conflict, and we suggest potential avenues for improving the resolution of the pancrustacean tree of life, especially by identifying the most crucial lineages to sample in future studies.

## Results

### Dataset

Our phylogenomic analyses (see Supplemental Methods) were conducted in an iterative approach, gradually expanding taxon sampling and outgroups from a more Malacostraca-specific study to all Pancrustacea, but using the same methods for ortholog selection and phylogenetic analysis. In total, we analyzed 3 different taxon sets (Datasets 1–3) (Table 1, S1–3). Dataset 3, the final iteration, had the largest taxon set, the most thorough phylogenetic investigations, and the most robust results, so most of this study focuses on the results of those analyses (Figs. 2, 4; Table S1, S4). However, through our investigation of these 3 varying taxon sets, we identified several reasons for topological differences among analyses and potential sources of error to be wary of, so we briefly summarize our iterative analyses here. Our first iteration, Dataset 1, comprised 23 malacostracans and a single thecostracan as outgroup (Table 1, S3). In ASTRAL analyses, despite sampling 7,652 gene trees, we recovered a paraphyletic Malacostraca with Thecostraca sister to Amphipoda, an obvious artifact likely due to long branch attraction (LBA) (Fig. S4). This led us to greatly expand taxon sampling for Datasets 2 and 3, including all major pancrustacean clades with chelicerate and myriapod outgroups collectively comprising 98 and 105 taxa, respectively (see Table S1 and Fig. 2 for Dataset 3, and Table S2 and Fig. S5 for Dataset 2), which we then interrogated in detail with additional phylogenetic methods (partitioned ML, partitioned ML with Dayhoff recoding, ML with C60 mixture models, PhyloBayes CAT-GTR, and ASTRAL). Even in the context of expanded taxon sampling, outgroup selection, and phylogenetic tree search methodology, we still recovered suspect relationships in Dataset 2, particularly that in all analyses we recovered monophyletic Xenocarida as sister to all other allotriocaridans, and Copepoda as sister to Hexapoda (Fig. S5). Given these surprising findings, we again interrogated our taxon selection and added 13 additional hexapods, 3 stomatopods, 2 remipedes, 2 copepods, 2 barnacles, and another leptostracan. In order to better balance the taxon sampling, we also removed taxa that were closely related to other species already sampled, which included: 8 amphipods, 3 decapods, 3 isopods, and the second hymenopteran; this resulted in Dataset 3 (Fig. 2, Table 2, Table S1.).

**Table 1.** Comparison of Datasets 1–3 assembled in this study

| Dataset | num.<br>taxa | num.<br>"crustacean"<br>orders | IG taxa | OG taxa | orthologs | alignment<br>length (aa) |
| --- | --- | --- | --- | --- | --- | --- |
| Data set 1 | 25 | 9 | Malacostraca | Thecostraca | 7,652 | 4,553,140 |
| Data set 2 | 98 | 29 | Pancrustacea | Chelicerata, Myriapoda | 559 | 80,215 |
| Data set 3 | 106 | 29 | Pancrustacea | Chelicerata, Myriapoda | 576 | 121,508 |

**Table 2.** Comparison of recent pancrustacean phylogenomic analyses.

| Study | Number<br>"crustacean" orders | Number<br>orthologs | AA<br>positions | gaps per<br>ortholog | % gaps in<br>super matrix | Orthology<br>inference |
| --- | --- | --- | --- | --- | --- | --- |
| This study | 29 | 576 | 121,508 | 10% | 47% | tree-based |
| von Reumont (2012) | 14 | 316 | 62,638 | NA | 38% | sequence<br>similarity |
| Oakley et al. (2013) | 22 | 1,002 | 263,306 | NA | 80% | sequence<br>similarity |
| Lozano-Fernandez et al. (2016) | 6 | 246 | 40,657 | NA | 36% | tree-based |
| Schwentner et al. (2017) | 19 | 1077 | 301,748 | NA | 34% | sequence<br>similarity |
| Schwentner et al. (2018) | 25 | 455 | 112,993 | NA | 38% | sequence<br>similarity |
| Lozano-Fernandez et al. (2019) Matrix A | 23 | 244 | 57,149 | 25% | 25% | tree-based |
| Lozano-Fernandez et al. (2019) Matrix B | 23 | 2,718 | 53,039 | 23% | 28% | sequence<br>similarity |

**Figure 2.**
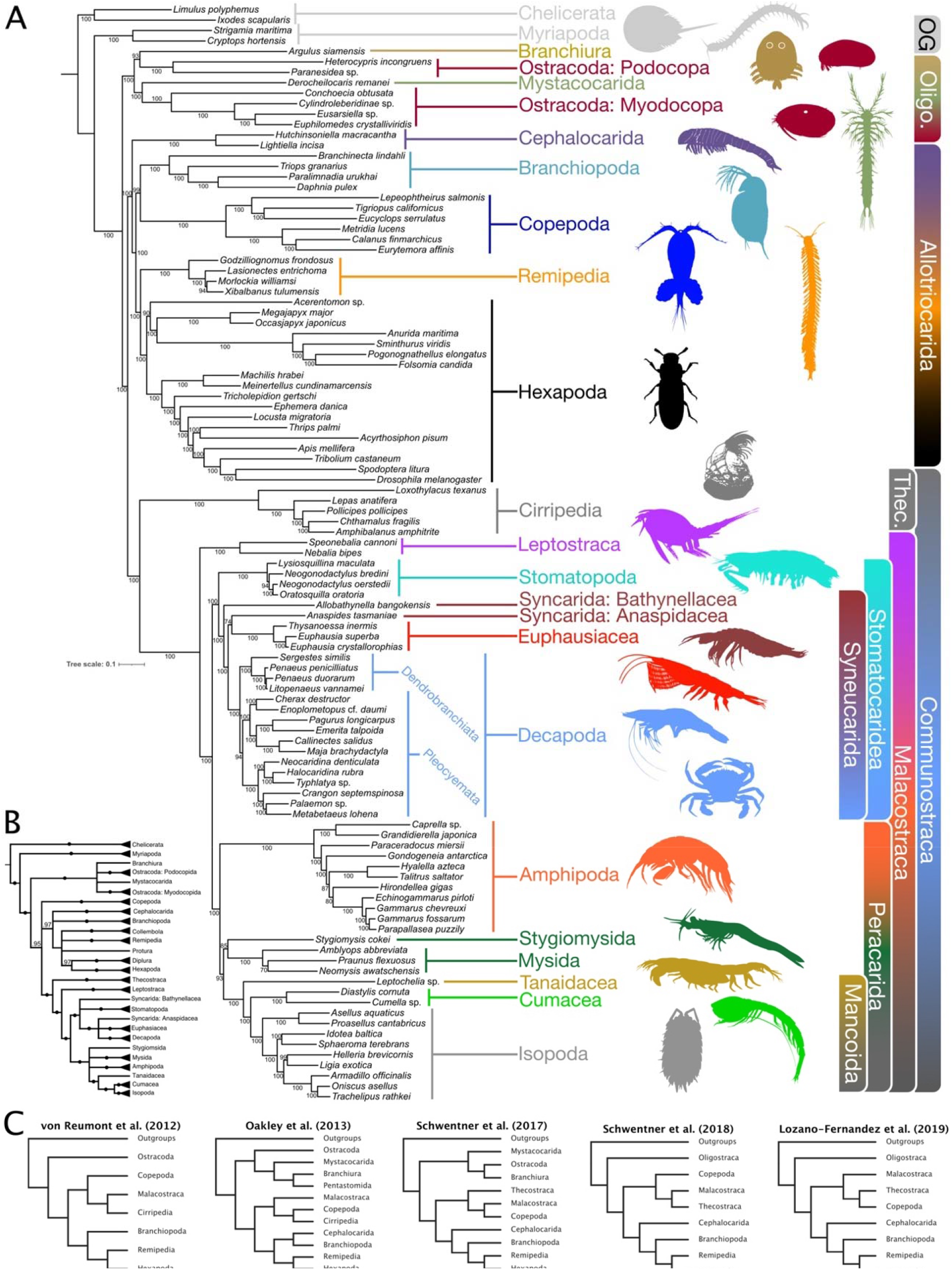
(A) Tree resulting from Maximum Likelihood analysis of the full matrix using LG+C60+F+G model. (B) Tree resulting from ASTRAL analysis of all orthologs with branches in gene trees with < 30% BS support collapsed, PP support values are 100% unless otherwise indicated. (C) Recent phylogenomic hypotheses of pancrustacean relationships.

Dataset 3, our final taxon set, is the primary focus of the study. We used genomic and transcriptomic data representing 29 of 55 crustacean orders and a phylogenetic diversity of hexapods. In total, 93 transcriptomes and 12 genomes spanning the arthropod tree of life were used for this study (Table S1). We had particularly high representation of malacostracans not sampled in previous phylogenomic analyses. We identified 576 protein-coding genes with a taxon occupancy >50% with an ortholog occupancy of 84–533 (average = 363, 63%) per species and a taxon occupancy of 53–99 (average = 66, 62%) species per ortholog. The taxon and ortholog occupancy statistics are summarized in Table S1; statistics for each of the orthologs are given in Table S4. The final concatenated alignment consisted of 576 orthologs, 121,508 amino acid positions, and 91,467 parsimony informative sites.

### Taxon Sampling Results

Somewhat surprisingly, with the relatively small changes in taxon sampling between Datasets 2 and 3 (75% of taxa shared) (Table 1), we found substantial differences in the topology of all ML, CAT-GTR, and ASTRAL analyses (Fig. 2, Fig. S5). Given that identical methods were used to call orthologs and for phylogenetic reconstruction, we reasoned taxon sampling was driving the differences in topology. Taxon sampling can generally affect 2 parts of a phylogenomic analysis: first, the ortholog selection (because clustering algorithms are sensitive to orthogroup structure, which is impacted by phylogenetic relatedness of taxa [Chen et al. 2007; Altenhoff and Dessimoz 2009]), and second, the accuracy of the reconstructed tree topology. We sought to identify through which of these processes taxon sampling was having its effects. To investigate this, we controlled for the effect of taxon sampling on ortholog selection by using only those orthologs that were exclusively shared between the two different taxon data sets; that is, we used the same subset of genes for phylogenetic tree searches of both taxon sets. Approximately ½ of orthologs were shared between the two datasets (267 of 559 and 576 orthologs, in Datasets 2 and 3 respectively). Using these same genes and the same ML, BI, and coalescent methods, we again recovered very different topologies between these 2 taxon sets. In fact, the topologies were nearly identical to the original topologies recovered from analysis of the full matrices from Dataset 2 and 3 (i.e., all 559 and 576 orthologs, respectively).

The Long Branch (LB) scores between Dataset 2 and 3 suggest that longest branches in the phylogeny are attributed to the Cirripedia, Copepoda, Ostracoda: Podocopa, Hexapoda, Branchiopoda, Ostracoda: Myodocopa, and the outgroup. In the Dataset 2 phylogeny, the LB scores ranged from −29.3-50.3, while in the Dataset 3 phylogeny they ranged from −28.3-37.1 (Table S5). In Dataset 2, the top 10% of the largest LB scores included taxa within Cirripedia (*Loxothylacus texanus*), Copepoda (*Lepeophtheirus salmonis, Tigriopus californicus, Eurytemora affinis*), Hexapoda (*Drosophila melanogaster, Folsomia candida*), Branchiopoda (*Brachinecta lindahli*), and Ostracoda: Myodocopa (*Conchoecia obtusata*). In Dataset 3, Branchiopoda, Ostracoda: Podocopa, and Ostracoda: Myodocopa were not in the top 10%, while the composition of Copepoda and Hexapoda taxa within the top 10% LB scores changed from Dataset to 3. The rest of our study focuses on our final taxon set, Dataset 3.

### Phylogenetic Results

We completed phylogenetic analysis using the following methods: partitioned ML analyses with RAxML, site heterogenous ML analyses (C60 family of models) with IQTREE2, Bayesian Inference site heterogenous CAT-GTR with PhyloBayes, and coalescent analyses with ASTRAL-III. While some topological differences occurred across the BI, ML, and coalescent-based methods, there was general agreement across analyses, especially among ML analyses (Fig. 3, Figs. S1–3). All methods supported the following topological arrangements: Oligostraca is the first group of pancrustaceans to diverge from all the others and contains a polyphyletic Ostracoda; within Altocrustacea (i.e., all pancrustaceans except Oligostraca), Thecostraca is the sister to the Malacostraca (Communostraca hypothesis of Regier et al. [2010]); Allotriocarida is the sister to Communostraca but is expanded to include Copepoda; Peracarida is monophyletic with Amphipoda in an early diverging position (Fig. 2A, B); and Decapoda and Euphausiacea form a clade with a paraphyletic Syncarida. The presence of the two clades of syncarids in this latter grouping, with Euphausiacea being closer to one of them (Anaspidacea) than to Decapoda, renders the classically recognized Eucarida polyphyletic and we propose the name Syneucarida for this expanded clade (Fig. 2A).

**Figure 3.**
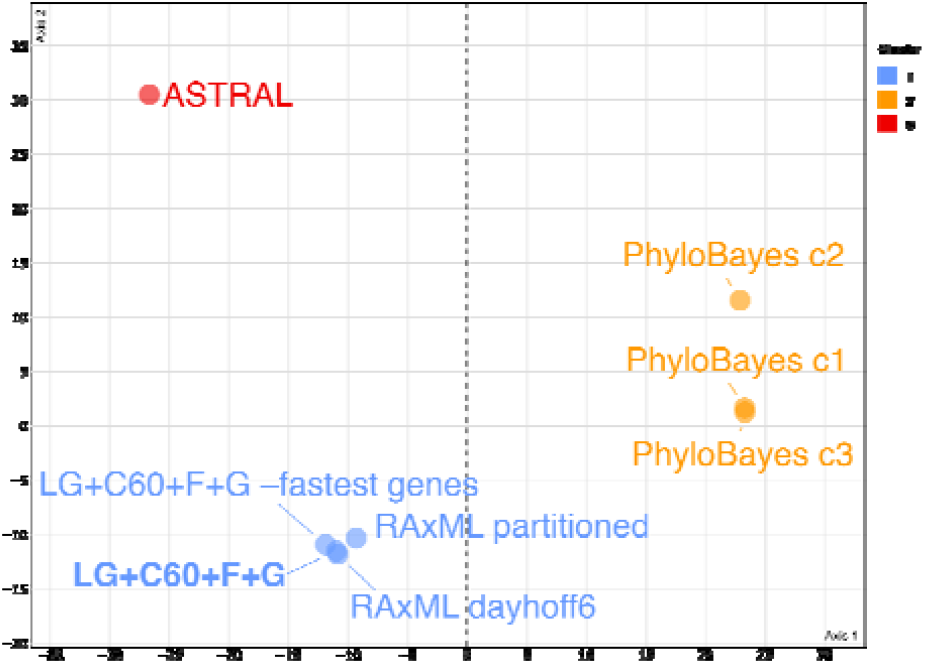
MDS of topological tree space of phylogenetic analyses in this study using the Kendall and Colijn (2016) method for defining summary trees.

While the ML, BI, and multispecies coalescent results showed a large degree of congruence, especially among ML analyses, the topologies differed in some parts of the pancrustacean tree (Fig. 3, Figs. S1–3). The tree resulting from the ML analysis using the LG+C60+F+G mixture model (Fig. 2A) is presented as our primary species tree for five reasons: (1) it was the substitution model of best fit by BIC with the caveat that the CAT-GTR model can not be tested with standard tests of model fit; (2) the AU-test (Shimodaira 2002) rejected the other topologies produced by other methods (i.e., ASTRAL and BI) in favor of this one (p < 0.001) (Table S6); (3) the 3 chains of the PhyloBayes CAT-GTR analysis did not fully converge (maxdiff between chains 0.69–1) even after approximately one year of continuous run time; (4) site-heterogeneous models like the C60 class account for among site variation in amino-acid propensities are less prone to artifacts like long branch attraction (Lartillot and Philippe 2004; Lartillot et al. 2007; Le et al. 2008) and were found to converge in this dataset in a reasonable time frame; and (5) there was high congruence between this topology and those produced from other methods, including a nearly identical topology from the partitioned RAxML analysis with and without Dayhoff recoding (Fig. 3). The resulting LG+C60+F+G phylogeny is the focus of most of this study, but differences between this topology and those from the other analyses are reviewed below.

Despite general agreement, the majority-rule posterior consensus tree of the CAT-GTR chains differed from the ML topologies in a few ways. The phylogenetic position of Stomatopoda was consistent in all ML and ASTRAL analysis, but differed under the CAT-GTR model: all three chains recovered Stomatopoda as the sister taxon to all other malacostracans except for the Leptostraca (Fig. S2). Still, given the consistency of the more derived position for Stomatopoda in ML and coalescent methods, we propose the name Stomatocaridea for referring to the new clade comprising Stomatopoda, Decapoda, and Syneucarida; this divides Malacostraca into three clades: Leptostraca, Stomatocaridea, and Peracarida.

Relationships within Allotriocarida also differed between ML and CAT-GTR analyses. While all ML analyses found Copepoda + Branchiopoda as the sister to Remipedia + Hexapoda, results varied between CAT-GTR chains. One chain found Copepoda alone as the sister to Remipedia + Hexapoda, while the two other chains found Copepoda + Remipedia as the sister to Hexapoda. These two different positions for Copepoda are interesting but were not recovered in any ML or ASTRAL analyses. The only other differences among our ML and BI analyses were in shallow nodes within clades. The exact relationship among the 3 mysids: *Neomysis, Praunus*, and *Amblyops* varied, one of the few branches to vary among ML analyses, because *Praunus* and *Amblyops* had the lowest and third lowest number of orthologs. Within Amphipoda, *Gondogeniea* was the sister to *Hirondellia* in one ML analysis, when the top 10% fastest evolving genes were removed, but in all other ML, BI, and ASTRAL analyses *Gondogeniea* was the sister to the large *Hyallela* + *Gammarus* clade.

ASTRAL topologies were more different from ML than BI (Fig. 3A, B, Fig. S1–3). Given that short exons (e.g., average of 211 AA here) are the predominate orthologs in arthropods that are stable over their >500 million year evolutionary history (e.g., Owen et al. 2020), ASTRAL may have suffered from relatively weak signal in individual gene trees or high gene tree error. As noted by others, summary methods like ASTRAL can be inappropriate when gene tree estimation error is high (Huang et al. 2010; Bayzid and Warnow 2013; Patel et al. 2013; DeGiorgio and Degnan 2014; Mirarab et al. 2014; Lanier and Knowles 2015; Mirarab and Warnow 2015; Xi et al. 2015). As a result, we interpreted results from ASTRAL, especially those that conflicted with ML or BI, with some skepticism. With that in mind, we summarize the main areas of conflict below. Within Allotriocarida, ASTRAL topologies showed a lack of congruence with other methods. ASTRAL consistently recovered Copepoda as the sister to all other Allotriocarida, while ML and BI methods consistently had Cephalocarida in this early diverging position (Supplemental Fig. S3A–C). Strikingly, ASTRAL also recovered Hexapoda paraphyletic with Protura + Diplura + Insecta more closely related to Remipedia than to Collembola (Supplemental Fig. S3) albeit with low support. This is almost certainly an artifact. In the full ASTRAL analysis, the node leading to Remipedia sister to Protura + Diplura + Insecta has 0.96 PP, but support for this node decreased when nodes with low support in gene trees were collapsed to polytomies (Supplemental Fig. S3A). When gene tree nodes with <10% and <30% BS support were collapsed prior to ASTRAL, support for the Remipedia + Protura + Diplura + Insecta node decreased to 0.91 PP and 0.45 PP respectively, demonstrating that this node in ASTRAL was supported by gene trees with low support at this node (Supplemental Fig. S3B, C). Another notable difference in ASTRAL relates to the evolutionary relationships of the mysids, but instability in this group is likely due to their low ortholog occupancy (*Praunus, Amblyops*, and *Stygiomysis* ranked first, third, and fifth in fewest orthologs of all taxa) (Table S1). Contrary to ML and BI analyses of the full amino acid matrix, ASTRAL and ML analysis of the Dayhoff6 recorded matrix found Mysida was paraphyletic with Stygiomysida as the sister to Mancoida and the other mysids sister to amphipods, but with low support (50% BS, <0.40 PP in ASTRAL). These results in ASTRAL and under Dayhoff recoding were treated with skepticism due to low number of orthologs and weak phylogenetic signal for *Praunus, Amblyops*, and *Stygiomysis* (Table S1).

### Divergence Time Estimation Results

We completed divergence time estimates across Pancrustacea using three chains each in MCMCTree and in PhyloBayes with autocorrelated (CIR and lognormal), and uncorrelated (UGAM) clock models. Fossil dates and justifications are given in Table S7. Convergence between chains was assessed by plotting posterior means for each chain against one another for MCMCTree (Fig. S7) and with trace plots for PhyloBayes (Fig. S8). Divergence time estimates from MCMCTree are summarized in Figure 4. Unlike the only previous study conducted with similar fossil calibrations (Schwentner et al. 2017), our use of MCMCtree allowed our age estimates to incorporate the full matrix. With these more extensive sequence data in MCMCtree, we retrieved deep root ages for arthropods, pancrustaceans, and the three major clades of pancrustaceans, extending slightly past and into the middle of the Ediacaran period for each. With our supplementary PhyloBayes analyses using only 50 loci (Fig. S10), these deeper nodes diverged within the Cambrian.

**Figure 4.**
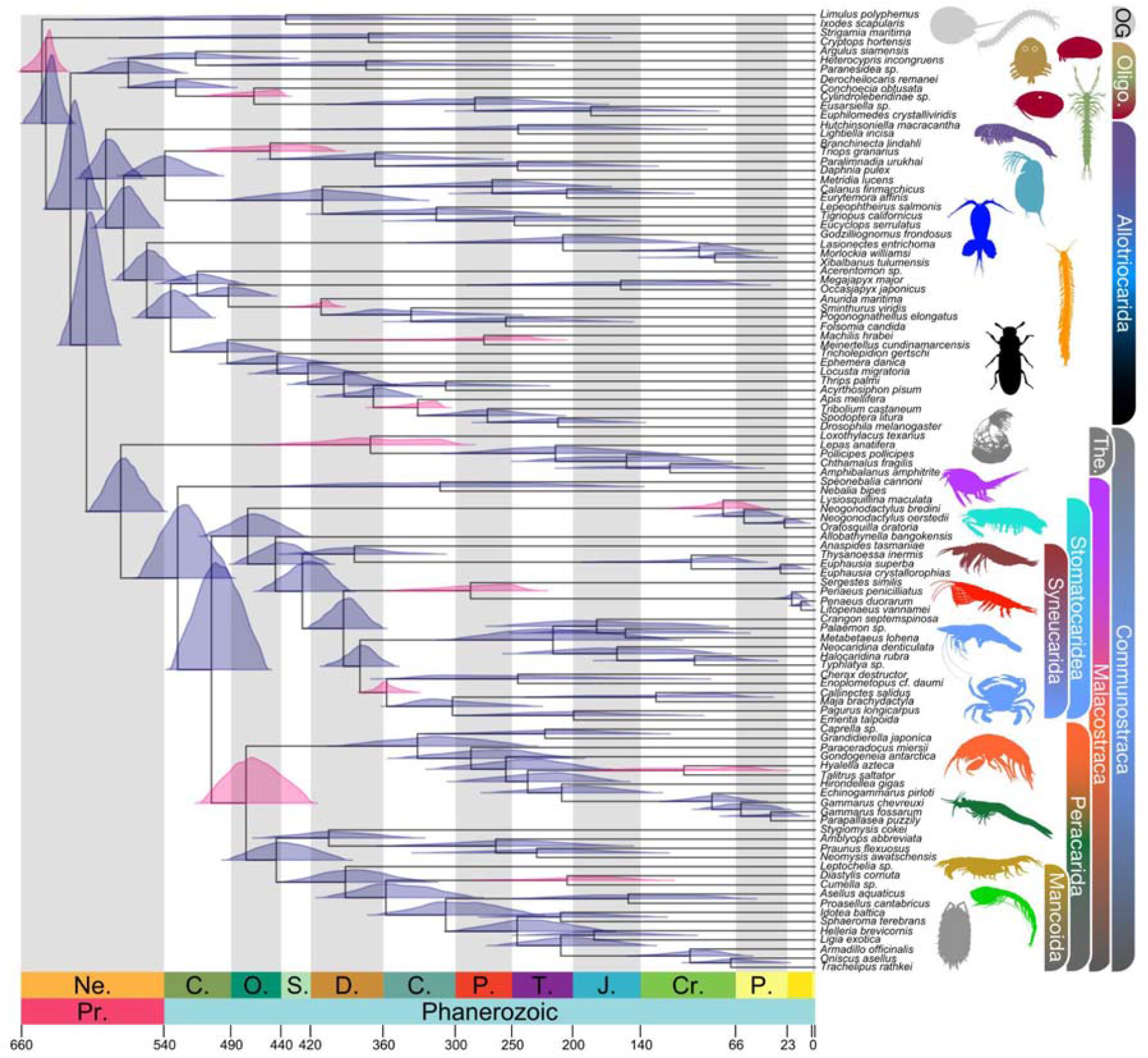
Fossil calibrated divergence time estimates for Pancrustacea, based on an MCMCtree analysis of the topology depicted in Figure 2A calibrated with 13 vetted fossils. The fossil calibrated nodes have their posterior age distributions highlighted in pink.

In the MCMCtree analysis, within Allotriocarida, hexapods were estimated to have terrestrialized in the late Cambrian. In all PhyloBayes models, terrestrial hexapods were estimated as Ordovician. Their sister group, remipedes, has a very wide 95% HPD with MCMCtree, reflecting a crown group that may have diverged in the Jurassic (mean age), with a range from Pennsylvanian to Cretaceous. The other large allotriocarid group without an internal calibration, copepods, likely diverged in the Devonian (with a range from Ordovician to end Permian depending on the clock model). Within Multicrustacea, major crown groups showed mean estimates for their divergences in the Ordovician (peracarids and syneucarids), Devonian (decapods), and Cretaceous (stomatopods). Finally, all sampled higher-level clades within Pancrustacea had diverged prior to the Cenozoic, a result that is consistent using PhyloBayes.

## Discussion

### Impacts of outgroup choice and taxon sampling

Our analyses took an iterative approach, gradually expanding taxon sampling and outgroup choice while using the same methods. In total we analyzed three different taxon sets, which yielded some insights into the causes of different topologies recovered in our analyses. Our first analysis had only 25 taxa: 24 malacostracans and a thecostracan as a putative outgroup, and produced some radically unusual relationships (e.g., paraphyletic Malacostraca with Thecostraca sister to Amphipoda) that we suspected were artifacts (Fig. S4). Our suspicions were confirmed by subsequent analyses with expanded taxon sampling and more rigorous phylogenetic methods that did not recover these highly unusual relationships. These results serve as a warning: despite sampling 7,652 orthologs in Dataset 1, we recovered potentially spurious relationships that appear to have been driven, at least partially, by limited outgroup selection (i.e., a single thecostracan as OG for Malacostraca [Figure S4]). Others have also found that limited outgroup selection can have a large impact on molecular analyses (Puslednik and Serb 2008; Rota-Stabelli et al. 2011), which should be considered in future studies. Our subsequent taxon iterations had greatly expanded taxon sampling and used chelicerate and myriapod outgroups. After recovering some unusual relationships in Dataset 2 (e.g., Copepoda sister to Hexapoda, Stomatopoda sister to syncarids and eucarids), we added additional hexapods, copepods, and stomatopods. We also removed some closely related amphipods, decapods, and isopods at the tips of the tree to better balance the taxon sampling such that malacostracans made up 50% of the dataset rather than 70%.

Because we recovered substantially different topologies between Datasets 2 and 3, which differed only in taxon sampling with 75% of taxa shared, we further investigated the effects of taxon sampling. We reasoned taxon sampling could be affecting two components of phylogenomic analysis: ortholog selection and tree topology accuracy. To disentangle these effects, we controlled for ortholog selection by using the same genes: only the 267 orthologs that were shared between the two datasets (roughly 50% of the orthologs). Using identical orthologs and only slightly different taxa, we repeated the same phylogenetic analyses and once again recovered incongruent topologies — topologies that were nearly identical to the originals recovered from analysis of the full ortholog alignments of each dataset (Figs. S5, S6). So while taxon sampling did impact ortholog identification (in that only half of orthologs were shared), by controlling for ortholog selection, our results demonstrate that the differences in topology between the two taxon sets was driven not by differences in the loci, but strictly on the phylogenetic reconstruction method, an impact we found surprisingly large given the small change in taxon coverage.

Although the results of the controlled ortholog experiment were surprising, we do find evidence that supports decades of literature suggesting taxon sampling and density increases phylogenetic accuracy. Arguably the biggest differences between Dataset 2 and Dataset 3 are the relationships estimated within Allotriocarida. In Dataset 2, we estimated a sister relationship between copepods and hexapods (Sup. Fig. 5), while in Dataset 3 we estimated the commonly supported sister relationship between remipedes and hexapods (Fig. 2). The taxon sampling differences between these datasets for these lineages are (1) an increase in the number of basal non-insect hexapods (i.e., Entognatha: Diplura, Collembola, and Protura) from one representative (Dataset 2) to seven (Dataset 3), (2) an increase in remipede taxa from two representatives (Dataset 2) to four (Dataset 3), and (3) an increase in copepods from four (Dataset 2) to six (Dataset 3). We believe that the reduced taxon sampling caused the spurious relationships in Dataset 2, due in part to the relative branch lengths of these groups. Generally speaking, the branches near the common ancestors of these lineages are quite short, but nearly all of the branches are relatively long towards the tips. Despite using a substitution model that accounts for non-stationarity and has been shown to be robust against LBA (Lartillot et al. 2007), we estimated copepods were the sister to hexapods in Dataset 2 with a single taxon representative of basal hexapods that had a branch length of 0.84 amino acids substitutions per site (Sup. Fig. 5). In Dataset 3, although there are still relatively long branches in the basal hexapods, the additional taxa decreased the average terminal branch length in this lineage from 0.84 to 0.27 amino acid substitutions per site. Moreover, we increased the taxon density for basal hexapods sevenfold, reducing the effects of LBA, recovered the traditional relationships, and ultimately confirmed that taxon sampling is important in the phylogenomics era. Other phylogenomic studies are also finding that subsampling deep lineages may cause topological inaccuracies (e.g., Branstetter et al. 2017; Betancur□R. et al. 2019) and the suspect relationships recovered with the limited backbone sampling in Dataset 1 further corroborates these conclusions.

### Comparison to other pancrustacean phylogenomic studies

In regard to taxon sampling, relative to previous studies, we sampled more crustacean orders (i.e., 29 vs <25) and sampled them more densely (e.g., 54 malacostracans vs 4–14) (Table 2). This was especially true within Malacostraca where we sampled more syncarids, peracarids (especially isopods and amphipods), decapods, and stomatopods than any prior study. We assembled our final matrix (Dataset 3) with an emphasis on maintaining balanced taxon sampling to the greatest extent possible. Taxon sampling was balanced by (1) selecting taxa representing deep splits in clades with known phylogenetic relationships and (2) when phylogenetic relationships were unclear, sampling as many taxa as possible followed by iteratively building species trees and subsampling clades to prune closely related species at the tips of densely sampled clades, retaining primarily the deep splits within clades. This was done because uneven taxon sampling is known to affect homolog clustering and therefore ortholog selection (Chen et al. 2007; Altenhoff and Dessimoz 2009), and because less balanced taxon sampling had a large impact on topology in our analyses of Dataset 2, even when controlling for ortholog selection (Fig. S6).

In general, most pancrustacean phylogenomic studies have shared a high degree of overlap in taxon selection and methodology. All have focused on single-copy protein-coding genes under one of three phases of data generation: Sanger sequencing (Regier et al. 2005; Regier et al. 2008; Regier et al. 2010; Rota-Stabelli, Lartillot, et al. 2013), expressed sequence tags (ESTs) from 454 sequencing (von Reumont et al. 2012; Oakley et al. 2013), and Illumina-based RNA-Seq (Schwentner et al. 2017; Schwentner et al. 2018; Lozano-Fernandez et al. 2019). Besides the Sanger sequence studies, most have had roughly similar alignment sizes, once number of orthologs, AA positions, and gaps are accounted for (Table 2). Despite these similarities, the topologies recovered have been surprisingly variable.

Beyond taxon sampling, our study differed from most others in several ways. First, we used a tree based approach to ortholog identification, which has been shown to improve phylogenetic reconstructions in simulation studies (Smith and Pease 2017) and published data sets (Dunn et al. 2013; Yang and Smith 2014; Ballesteros and Hormiga 2016). A few other pancrustacean studies have used a tree-based approach to ortholog identification, but most have used the sequence similarity approach in OMA (Altenhoff et al. 2011) (Table 2). Second, we de-emphasized results recovered only with n Dayhoff 6-state recoding, a form of data reduction that may remove true signal more than it ameliorates saturation and compositional heterogeneity (Hernandez and Ryan 2021) (though we do present a Dayhoff recoded ML tree [Fig. S1B and Fig. 3]); instead, we favored accounting for saturation and compositional heterogeneity with site-heterogenous models in addition to partitioned and coalescent analyses. Third, we did not rely primarily on results under the CAT-GTR model. In pancrustacean phylogenomic analyses, CAT-GTR chains frequently do not fully converge, as was the case here despite nearly a year of run time, and without convergence the results are statistically invalid (Gelman and Rubin 1992; Huelsenbeck et al. 2002; Whelan and Halanych 2017). Furthermore, Li et al. (2021) showed that CAT-GTR analyses of the metazoan tree of life often have hundreds of additional rate categories yet fail to fit better than site-heterogeneous models with many fewer categories. Given the issues with convergence under CAT-GTR, we emphasized the phylogeny resulting from the LG+C60+F+G mixture model, which has not been used in previous pancrustacean phylogenomic analyses. Site-heterogeneous models like the C60 class still account for among site variation in amino-acid propensities, are less prone to artifacts like long branch attraction (Lartillot and Philippe 2004; Lartillot et al. 2007; Le et al. 2008), and were found to converge in this dataset in a reasonable time frame. It was also the model of best fit, the highest likelihood tree, and was not rejected by the AU-test (unlike CAT-GTR) (Table S6). Finally, the tree resulting from the C60 analysis was robust; that is, it was nearly identical to those produced from the partitioned RAxML analysis with and without Dayhoff recoding (Fig. 3).

### Phylogeny and systematics

Some pancrustacean clades have been consistently recovered across phylogenomic analyses. The position and composition of Oligostraca has been relatively constant compared to the other major clades (Regier et al. 2010; von Reumont et al. 2012; Oakley et al. 2013; Rota-Stabelli, Lartillot, et al. 2013; Schwentner et al. 2017; Schwentner et al. 2018; Lozano-Fernandez et al. 2019). Although a few analyses have found Oligostraca to be more closely related to Malacostraca + Thecostraca (see von Reumont et al. [2012] figure 2 and Rota-Stabelli, Lartillot et al. [2013]), the vast majority have recovered Oligostraca as the sister to all other pancrustaceans (Regier et al. 2010; von Reumont et al. 2012; Oakley et al. 2013; Schwentner et al. 2017; Schwentner et al. 2018; Lozano-Fernandez et al. 2019). A major question that remains is the monophyly of ostracods. Oakley et al. (2013) recovered a monophyletic Ostracoda with relatively dense sampling of ostracods, but more recent studies that have relied primarily on ostracod transcriptomes rather than ESTs and have included fewer ostracods as a result, have often found a polyphyletic Ostracoda, as we have here. Clearly, expanded taxon sampling of the Ostracoda is needed, which will also enable the incorporation of the rich fossil record of ostracods for divergence time analyses. The question of ostracod monophyly will be best explored not just in the context of expanded sampling of ostracods (Ellis et al. 2022) but also by expansion of the poorly sampled Mystacocarida and Branchiura. In most recent phylogenomic studies, Mystacocarida is represented only by *Derocheilocaris remani* and Branchiura only by *Argulus siamensis* (von Reumont et al. 2012; Rota-Stabelli, Lartillot, et al. 2013; Schwentner et al. 2017; Schwentner et al. 2018; Lozano-Fernandez et al. 2019). Therefore the phylogeny of Oligostraca would benefit most from expanded taxon sampling of ostracods (especially *Manawa staceyi* and Platycopa), the other genus of Mystacocarida (*Ctenocheilocaris*), and other ichthyostracans; in the case of Ichthyostraca, sequencing the first pentastomid transcriptome and the early-diverging branchiuran *Dolops* would be particularly valuable (Møller et al. 2008).

Most pancrustacean phylogenomic studies have recovered a clade comprising some combination of Branchiopoda, Cephalocarida, Hexapoda, and Remipedia, in a group termed Allotriocarida. In terms of the interrelationships, the sister group to hexapods has received the most attention due to potential insights about pathways of terrestrialization. Regier et al. (2010) found Remipedia + Cephalocarida as the sister to Hexapoda but subsequent studies have usually found Remipedia alone sister to Hexapoda, with Cephalocarida diverging earliest from the rest of Allotriocarida (Oakley et al. 2013; Schwentner et al. 2017; Schwentner et al. 2018; Lozano-Fernandez et al. 2019). Multiple studies have noted the Cephalocarida + Remipedia pairing may be an artifact of LBA (Rota-Stabelli, Lartillot, et al. 2013; Schwentner et al. 2017; Lozano-Fernandez et al. 2019). We recovered Cephalocarida as the sister to all other allotriocaridans with maximum support in all concatenation-based analyses. Branchiopods have been relatively well-sampled by Schwentner et al. (2018) and remipedes by Lozano-Fernandez et al. (2019), but with only 2 cephalocarid transcriptomes to date, the phylogeny of Allotriocarida would benefit from sampling members of the 3 remaining cephalocarid genera (WoRMS 2022).

We recovered Allotriocarida with the marked addition of Copepoda in all analyses (i.e., ML, BI, and multispecies coalescent). The analyses in this study provide robust support for an Allotriocarida that would be expanded to include Copepoda, but the exact position of copepods is less clear. All of our ML analyses (partitioned, site heterogenous, Dayhoff recoded, and subsampled matrices) consistently found copepods sister to branchiopods with high support (95–100% BS). In Phylobayes, 2 CAT-GTR chains recovered Copepoda + Remipedia as the sister to Hexapoda with 100% PP, while the third chain found Copepoda sister to Remipedia + Hexapoda. Meanwhile all ASTRAL analyses estimated Copepoda sister to all other allotriocaridans (Figure S3), but with consistently low support, highlighting a lack of congruence among methods in this part of the phylogeny. Two prior studies occasionally recovered Copepoda in Allotriocarida in a subset of their analyses, but the position of copepods was variable: Lozano-Fernandez et al. (2019) found Copepoda sister to Remipedia (their figure 1B) or Remipedia + Hexapoda (their figure 1C), while Rota-Stabelli et al. (2013) found Copepoda sister to Branchiopoda (their figures 1C, D). There is some morphological support linking Copepoda to allotriocaridan taxa. Over 30 years ago, Ito (1989) was one of the first to hypothesize an evolutionary relationship between Copepoda, Remipedia, and Cephalocarida based on his observations of their limb morphology. He noted copepods and remipedes both possess six cephalic limbs and further proposed a scenario where the three-segmented endopod of the copepod trunk limb was derived from an ancestral remipede-like four-segmented endopod, a suggestion further supported by the fact that some remipedes already possess three-segmented endopods on their posterior trunk limbs. Additionally, Ito (1989) proposed that these three and four-segmented endopods may have been derived from an ancestral 5-segmented endopod like those seen in cephalocarid trunk limbs. Still, the exact position of copepods continues to be one of the least resolved parts of the pancrustacean tree of life (Rota-Stabelli et al. 2013; Lozano-Fernandez et al. 2019). We think a more precise position of Copepoda rests in sampling Platycopioda, the copepod order sister to all others (Huys and Boxshall 1991), with genome-scale data. Species of Platycopioda have scarcely been sequenced at all, but this taxon should shorten the long branch leading to Copepoda, which can ameliorate phylogenetic error (Hendy and Penny 1989).

Nonetheless, the presence of Copepoda within Allotriocarida does have implications for the evolution of the other constituent clades. First, given that copepods were almost certainly ancestrally marine and hyperbenthic (Huys and Boxshall 1991), their close relationship with branchiopods, hexapods, and remipedes provides additional support for the hypothesis that these clades were also ancestrally marine (von Reumont et al. 2012; Lozano-Fernandez et al. 2016; Schwentner et al. 2017). Second, Schwentner et al. (2017) noted that the loss of the mandibular palp in adults (though present in juveniles of Branchiopoda, Cephalocarida, and Remipedia) might be an apomorphy for Allotriocarida. However, the position of Copepoda recovered here (Fig. 2B) suggests the evolutionary history regarding the loss of the mandibular palp in adults is homoplasious. Either the mandibular palp was lost separately in the adults of Branchiopoda, Cephalocarida, and Remipedia, or Copepoda is unique in Allotriocarida for retaining the mandibular palp in adulthood; the latter scenario is more parsimonious and is supported by other neotenic features proposed for Copepoda (Gurney 1942).

Malacostraca and Thecostraca have a consistent phylogenetic affinity in all recent phylogenomic studies, but Multicrustacea (Copepoda, Malacostraca, and Thecostraca) has been one of the least stable areas of the pancrustacean tree of life, mostly due to variability in position of Copepoda (Regier et al. 2010; von Reumont et al. 2012; Oakley et al. 2013; Rota-Stabelli, Lartillot, et al. 2013; Schwentner et al. 2017; Schwentner et al. 2018; Lozano-Fernandez et al. 2019). Previous studies have typically found Copepoda as the sister to Malacostraca + Thecostraca, or sister to one of those taxa individually.

However, as noted above, some analyses, including all tree searches in this study, recover Copepoda within Allotriocarida, rejecting the traditional Multicrustacea (Copepoda + Malacostraca + Thecostraca). Instead, we recovered Communostraca (Malacostraca + Thecostraca) in all analyses. While Lozano-Fernandez et al. (2019) did not find consistent support for Communostraca across their analyses, our results support their suggestion that the location of male and female gonopores on different body somites is a synapomorphy for Communostraca. To further resolve the communostracan phylogeny and more comprehensively test the validity of “Multicrustacea”, it is important to sample Platycopioda and the early diverging thecostracan lineages Acrothoracida, Ascothoracida, Facetotecta, and Tantulocarida (Petrunina et al. 2014; Chan et al. 2021).

With 43,519 extant species (WoRMS 2022), Malacostraca is the most speciose crustacean class, but it has received relatively little attention in pancrustacean phylogenomic studies. Most did not comment on malacostracan interrelationships because they sampled only 4–13 species (Regier et al. 2010; von Reumont et al. 2012; Oakley et al. 2013; Schwentner et al. 2017). One constant is Leptostraca as the sister to all other malacostracans. Schwentner et al. (2018) included the most malacostracans prior to this study, and examined interrelationships among the 26 species sampled there, of which most (60%) were decapods. Here, we examined malacostracan relationships with expanded taxon sampling, doubling the number of species sampled (52), particularly peracarids and stomatopods. An earlier version of this analysis (Dataset 2) included 14 additional amphipods, isopods, and decapods, but these branches comprised shallow splits and these taxa were removed to maintain a more balanced taxon set. Our results regarding Malacostraca differ from those of Schwentner et al. (2018) in several ways. With the inclusion of Bathynellacea for the first time, we were able to test the monophyly of Syncarida (Anaspidacea + Bathynellacea) and it was recovered as polyphyletic in all analyses, with Anaspidacea as sister to Euphausiacea resulting in a paraphyletic Eucarida (Euphausiacea + Decapoda). Our results support the hypothesis of Serban (1972, 1973) that the Syncarida is polyphyletic with Bathynellacea in an earlier diverging position. Notwithstanding the non-monophyly of Syncarida and Eucarida respectively, a clade with all their subtaxa (Anaspidacea, Bathynellacea, Eucarida, and Decapoda) (Fig. 2), was recovered with maximum support in all ML and BI analyses for which we propose the name Syneucarida (Fig. 1A).

The position of Stomatopoda has been variable. While the CAT-GTR analysis in Schwentner et al. (2018) found Stomatopoda in a more classical, early diverging position in Malacostraca, all of our ML and ASTRAL analyses recovered Stomatopoda as sister to Syneucarida. We propose the name Stomatocaridea for Stomatopoda + Syneucarida, dividing Malacostraca into 3 clades: Leptostraca, Stomatocaridea, and Peracarida. Interestingly, our three CAT-GTR chains recovered Stomatopoda in the more basal position found by Schwentner et al. (2018), but we treated this result with some skepticism given that all ML and ASTRAL analyses consistently grouped Stomatopoda with Syneucarida (Fig. 2A, B), and that topology tests rejected the earlier diverging arrangement (Table 2). In the CAT-GTR analysis of the Dayhoff recoded matrix in Lozano-Fernandez et al. (2019), Stomatopoda was recovered as the sister to Mysida; we suspect this to be an artifact, perhaps due to LBA, given that we never recovered this grouping in any analysis under our expanded sampling of mysids and stomatopods. Several improvements can be made to better resolve relationships among stomatocaridean taxa (Stomatopoda and Syneucarida). Sampling more stomatopods may address the variability in their phylogenetic relationships; we attempted to include *Hemisquilla californiensis* (SRR2103462–3) and *Pseudosquilla ciliata* (SRR2103518, SRR2103524) in our study, but removed them from phylogenetic analyses because we were never able to recover more than 13% of orthologs from these samples. Species of *Hemisquilla* would be particularly valuable given a recent study suggested they are sister to the other stomatopods (Koga and Rouse 2021). Since both syncarid orders are represented by a single species each, sampling additional syncarids would be beneficial.

Our expanded taxon sampling of peracarids enabled us to examine relationships in this clade in greater detail than previous phylogenomic studies. In all ML and BI analyses, we found a monophyletic Peracarida with amphipods sister to all other peracarids and Mysida sister to Mancoida (Isopoda + Cumacea + Tanaidacea) (Fig. 2). Our results contrast Schwentner et al. (2018) in this respect, where results varied among methods and where Mysida was frequently sister to the other peracarids. With expanded taxon sampling here comprising two additional mysids, a stygiomysid, a second cumacean, eight and nine select isopods and amphipods respectively, we consistently recovered mysids sister to Mancoida in all concatenated analyses. Within Mancoida, all of our analyses grouped Cumacea sister to Isopoda, contrary to the Tanaidacea + Cumacea relationship recovered in most analyses in Schwentner et al. (2018). Our results also disagree with the morphological phylogenetic hypothesis of Richter and Scholtz (2001) that linked Isopoda + Tanaidacea. Interestingly, in ML analyses of a Dataset 2, which that included 8 more amphipods and 3 more isopods (all shallow branches that were pruned to create a more balanced taxon set here), we did find Mysida sister to all other peracarids, similar to Schwentner et al. (2018). This result, however, was not robust within that dataset: ASTRAL and CAT-GTR analyses of the same matrix consistently found mysids sister to Mancoida, just as in all concatenation methods of our final matrix here. Taken together, these differences among our analyses, as well as those of Schwentner et al. (2018) and Höpel et al. (2022), suggest a surprising amount of instability at the base of Peracarida. Peracarida is one of the pancrustacean taxa most in need of sampling effort. There are 12 extant orders of peracarids, and half of them have yet to be sampled in phylogenomic analyses (WoRMS 2022).

Resolving the backbone of the peracarid phylogeny requires sampling the six remaining orders: Bochusacea, Ingolfiellida, Lophogastrida, Mictacea, Spelaeogriphacea, and Thermosbaenacea. These orders are crucial not just for the peracarid tree of life, but also for the larger Malacostraca, especially given that some have questioned whether Lophogastrida and Thermosbaenacea belong in Peracarida at all (Siewing 1956; Schram and Hof 1998). A recent mitochondrial genome study by Höpel et al. (2022) included Lophogastrida and recovered a monophyletic Peracarida with Lophogastrida sister to Mysida and Stygiomysida, a hypothesis that would be interesting to test with nuclear loci. Sampling these taxa would enable a robust test of the monophyly of Peracarida and might provide more resolution for the position of Stomatopoda given the short branches found separating Peracarida, Stomatopoda, and Syneucarida here.

### Divergence time estimation

Our MCMCtree divergence time estimates retrieved deep splits of arthropods and the three main pancrustacean clades (Oligostraca, Allotriocarida, and Communostraca) earlier than the Cambrian, heavily contradicting the crown group arthropod fossil record (first appearing around 521 mya; Daley et al. 2018) and the pancrustacean fossil record (stem and crown groups simultaneously appearing about 514 mya; Zhai et al. 2019; Hegna et al. 2020). It has been proposed that molecular clock models may overestimate the time of divergence of the crown group MRCA, and that their stem groups may go extinct quickly after the MRCA, together suggesting a “long fuse” divergence estimate is unlikely (Budd and Mann 2020a). It is possible that our MCMCtree results, using the uncorrelated independent rates clock model, represent such an example of overestimation of the crown group age, as older root ages have been observed before with this software and clock model (Barba-Montoya et al. 2017). To further investigate, we compared MCMCtree to divergence times estimated under three different clock models in PhyloBayes (Figure S10). We found that the arthropod root was within the Cambrian using autocorrelated clock models (CIR and lognormal), and in the uncorrelated (UGAM) analysis crown group Pancrustacea diverged in the Cambrian. These chronograms all estimated shorter branch lengths for the presumed early, rapid divergences, while our results with MCMCtree appear to “smooth” the rate of early evolution at deep nodes. Another hint comes from the relatively narrow posterior age distributions at these deep nodes in all analyses (compared to the marginal priors; Figure S9, S10), with wide distributions at many shallow nodes, which suggest a decrease in rates over time that may challenge clock models (dos Reis, Thawornwattana, et al. 2015). It is not so simple as to assume that uncorrelated clock models overestimate divergence times in our dataset, as the PhyloBayes UGAM analysis resulted in the youngest ages for many internal nodes. It is therefore unclear what drives the differences among clock models for internal node age estimates, although perhaps autocorrelated models are better able to cope with putative rapid divergences in the Cambrian (Lee et al. 2013; Daley et al. 2018; Budd and Mann 2020b) and a subsequent slowdown, similar to that proposed for one possible topology of chelicerates (Lozano-Fernandez et al. 2020).

Broadly, most major pancrustacean clades (Oligostraca, Allotriocarida, Communostraca, and most classes) were established in the early Paleozoic. The Late Cambrian origin for terrestrial Hexapoda estimated by MCMCtree is consistent with some recent studies (e.g., Lozano-Fernandez et al. 2016; Schwentner et al. 2017), while the much younger age (at least 100 mya younger, up to 330 mya) of the sister group, crown Remipedia, presents further challenges to the quest for identifying a stem group of either clade in the fossil record, as a genuine ghost lineage indicates a long time to pioneer different habitats and many unknown morphological changes.

Most shallower posterior age estimates were roughly consistent with their fossil records (e.g., Wolfe et al. 2016; Hegna et al. 2020), highlighting the importance of appropriately vetted age priors in divergence time studies. Our new topological result supporting Syneucarida may prompt re-evaluation of Paleozoic fossils previously assigned to the extinct ‘syncarid’ group Palaeocaridacea (Schram 1984; Hegna et al. 2020). Although we were not able to include many peracarid fossil calibrations due to their lack of phylogenetic framework (Hegna et al. 2020), ages within the group were consistent in several cases, including isopods with a mean age range from the Carboniferous to Permian, depending on the clock model. Other clades which do not have fossil calibrations available (e.g., remipedes) or that were not appropriate to use with our particular molecular taxon sampling (e.g., copepods, most of amphipods) exemplified wider posterior ages under all clock models, often varying between models. In one standout case, penaeid shrimp, there are known crown group fossils from the Triassic (Wolfe et al. 2019), but they could not be used as priors and we retrieved impossibly young posterior ages with all clock models.

## Conclusions

Given that Pancrustacea contains >1,000,000 described species, a strategic expansion taxon sampling is required for better resolution of their phylogeny. The primary objective should be to sample the 26 crustacean orders that have not yet been sampled with genomic or transcriptomic data. Of utmost priority, we identify just 15 crucial taxa that likely represent deep branches in the pancrustacean tree; we suspect the inclusion of these few taxa may greatly improve the resolution of the pancrustacean phylogeny: *Manawa* and Platycopa (Ostracoda); Pentastomida and *Dolops* (Ichthyostraca); Platycopioida (Copepoda); Acrothoracida, Ascothoracida, Facetotecta, and Tantulocarida (Thecostraca); and Bochusacea, Ingolfiellida, Lophogastrida, Mictacea, Spelaeogriphacea, and Thermosbaenacea (Peracarida). Such taxa should be prioritized in genome sequencing efforts (Lewin et al. 2022). Until we have full genome sequences for the major branches of the tree of life (Lewin et al. 2018), we believe adding carefully curated sequences for taxa that have yet to be sampled is the most promising avenue for resolving the pancrustacean phylogeny.

In addition to expanding taxon sampling, greater resolution of the pancrustacean tree of life may be achieved by improvements in sequence data. Increasing the length and number of orthologs would add sites that could help resolve challenging nodes. Genomic sequences would be particularly valuable in this regard; of those crustaceans that have been sampled in phylogenomic analyses more than 90% are represented only by transcriptomes. High quality genome assemblies can provide more contiguous sequence data as well as non-coding sequences (e.g., Bernot et al. 2022). While identifying more orthologs is a worthwhile endeavor, such work must be done carefully and not only for the sake of more genes; the inclusion of a small number of paralogs can introduce strong erroneous signal that can mislead phylogenetic reconstructions (Shen et al. 2017; Smith and Hahn 2021), which underscores the importance of curation of the sequence alignments (Smith and Hahn 2022). It is noteworthy that the number of orthologs identified has been relatively consistent despite a variety of ortholog identification methods (Table 2), which may indicate that there are not many additional conserved orthologs across pancrustaceans to be added. Other methods in addition to tree-based orthology inference could help screen orthologs and reduce sources of error and improve phylogenetic reconstruction. Finally, more clade-specific analyses often yield more orthologs and more complete matrices, which may be useful for resolving particular areas of the pancrustacean tree (Schwentner et al. 2018; Laumer et al. 2019; Wolfe et al. 2019).

Additional data types could help resolve difficult areas of the phylogeny. New methods that do not rely solely only orthologs but also incorporate phylogenetic signal in paralogs, thus leverage substantially more data, may help resolve the pancrustacean phylogeny (Hellmuth et al. 2015; Smith and Hahn 2021; Smith et al. 2022). Phylogenetic signal can also be mined from synteny, which can be more conserved than coding sequence (Moret and Warnow 2005; Hu et al. 2014; Simakov et al. 2022).

Unfortunately, given the small number of chromosome scale crustacean genome assemblies (Bernot et al. 2022), probing phylogenetic signal from synteny in pancrustaceans is dependent on concerted genome sequencing efforts. Finally, in light of the recent discoveries of a number of incredibly preserved pancrustacean fossils (Zhang et al. 2007; Wolfe et al. 2016; Luque and Gerken 2019; Zhai et al. 2019; Robin et al. 2021) it is exciting to consider that new fossil discoveries may inform our understanding of the tempo, order of character acquisition, and patterns of diversity through time in pancrustacean evolution.

## Methods

### RNA extraction and sequencing

We collected fresh specimens of *Amblyops abbreviata, Diastylis cornuta*, and *Echinogammarus pirloti* and stored them in RNAlater (Invitrogen, Waltham, MA) or in no preservative at −80□. Total RNA was extracted using TRIzol (Invitrogen, Waltham, MA) according to the manufacturer’s instructions. Following extraction, total RNA was cleaned using the Nucleospin RNA Clean-up (Macherey-Nagel, Düren, Germany) silica-based column to further purify the RNA. Ribosomal RNA was removed using Ribo-zero (Illumina, San Diego, CA) and the quality of the RNA was checked on a Bioanalyzer. Both *Diastylis cornuta*, and *Echinogammarus pirloti* were sequenced on an Illumina HiSeq 2500 with 125bp paired-end reads at Duke University, while *Amblyops abbreviata* was sequenced on an Ion Torrent at the University of Bergen.

### Data set and transcriptome assembly

In total, 149 transcriptomes and 16 genome assemblies spanning the arthropod tree of life were used in this study (Supplemental Table S1–S3). Taxon sampling was iteratively expanded between Datasets 1, 2, and 3. The final taxon set, Dataset 3, (Table S1) comprised 106 transcriptomes and 14; 14 transcriptomes were subsequently removed from downstream analyses because they contained <20% of orthologs. The genomes of two chelicerates, *Limulus polyphemus* and *Ixodes scapularis*, and two myriapods, *Cryptops hortensis* and *Strigamia maritima*, were selected as outgroup taxa based on previous phylogenetic studies (Regier et al. 2010; Schwentner et al. 2017; Schwentner et al. 2018; Lozano-Fernandez et al. 2019). All computational analyses were carried out on the high-performance computing cluster at George Washington University.

Raw reads for all transcriptomes were assembled de novo as follows. Raw read quality was assessed using FastQC v0.11.8 (Andrews 2018), reads were subjected to quality and adapter trimming using Trimmomatic v0.33 (ILLUMINACLIP: TruSeq3-PE-2.fa:2:30:10 LEADING:3 TRAILING:3 SLIDINGWINDOW:4:15 MINLEN:50) (Bolger et al. 2014), and quality trimming and adapter removal was confirmed using FastQC again after trimming. Trimmed reads were error-corrected using Rcorrector (Song and Florea 2015) with default settings. Error corrected reads were assembled using Trinity (Grabherr et al. 2011; Haas et al. 2013) under default parameters. Assembled contigs were translated to amino acid sequences using TransDecoder v5.2.0 (Haas et al. 2013) with open reading frames identified using default parameters. Redundancy in amino acid sequences resulting from Transcoder was reduced using CDHIT v4.6 (Li and Godzik 2006; Fu et al. 2012) with a 99% similarity threshold.

### Ortholog identification

Orthologs were identified using an explicit phylogenetic approach following Yang and Smith (Yang and Smith 2014) (unless otherwise noted, named scripts are from https://bitbucket.org/yangya/phylogenomic_dataset_construction/src/master). The predicted proteins from the transcriptomes and genomes were subjected to an all-by-all BLASTP v2.9.0 (Altschul et al. 1990; Altschul et al. 1997; Camacho et al. 2009) search (-max_target_seqs 1000 -evalue 10) and the resulting BLAST output was filtered for the hit fraction being at least 0.4 (Chiu et al. 2006). Filtered BLAST hits were further clustered using MCL v12.068 (Van Dongen 2000; Van Dongen 2008) with a −log E-value cutoff set to 5 and an I-value of 1.4 to identify homologous protein sequences. Fasta files were written from the MCL output using write_fasta_files_from_mcl.py.

Each cluster of homologs was then aligned individually with MAFFT v7.13 (–genafpair– maxiterate 1000 if <1,000 sequences; –auto if >1,000 sequences) (Katoh and Standley 2013), trimmed using phyutility (minimum column occupancy = 0.1) (Smith and Dunn 2008), and trees were built using either RAxML v8.2.9 (Stamatakis 2014) under the model “PROTGAMMALG” for clusters with less than 1,000 sequences, or FastTree v2.1.8 (Price et al. 2010) under the model “-lg” for clusters greater than 1,000 sequences since the LG matrix was the model of best fit for the majority of orthogroups. The resulting trees may contain branches representing paralogs or misassembled contigs. These were identified and filtered using the following three methods. First, divergent sequences were removed from clusters if a terminal branch was longer than 1.0 or more than 15x longer than its sister using trim_tips.py. Next, if monophyletic or paraphyletic tips from the same taxa were present in a tree, only the sequence with the highest number of non-ambiguous characters in the trimmed alignment was kept and the rest removed following previously published methods (Smith et al. 2011; Dunn et al. 2013; Yang and Smith 2014). Lastly, potential deep paralogs were removed using cut_long_internal_branches.py with an internal branch length cutoff of 1.8 and a minimum number of taxa of 15. Fasta files were written from the trimmed trees and alignments and the entire process of aligning, trimming alignments, building trees, and removing paralogs and long branches was repeated. After the second round of refinement, the trees were called homolog trees and were further pruned to infer orthologs.

Orthologs were called using the maximum inclusion method (Dunn et al. 2008; Dunn et al. 2013; Yang and Smith 2014; Ballesteros and Hormiga 2016). After pruning the homolog trees to identify maximum inclusion orthologs, the remaining subtrees may contain terminal taxa subtended by long branches as a result of the subtree trimming method (Yang and Smith 2014). To account for this, the trees were trimmed once more using a range of permissive-to-strict branch length trimming parameters, referred to from here on as permissive, medium, and strict branch trimming, with relative branch lengths of 10x, 12x or 15x and absolute branch lengths of 0.75, 0.85, or 1.0 at the permissive, medium, and strict levels, respectively. Previous analyses showed that the more strict branch length trimming parameters only resulted in the loss of ~40 orthologs, so the strict branch length trimming was used here. The resulting orthologs were aligned with MAFFT and trimmed with Gblocks v0.91b using less strict parameters (Talavera and Castresana 2007) and the final matrix was produced using concatenate_matrices.py (Yang and Smith 2014) with a minimum number of sites set at 50 amino acids and a minimum taxon cutoff of 53/105 taxa (50.5%). The resulting matrix was composed of 576 orthologs.

### Phylogenetic analyses

Phylogenetic analyses were completed using concatenation and coalescent methods. Concatenation analyses were done in both maximum likelihood (ML) partitioned analysis, and ML and Bayesian inference (BI) mixture models. In the ML framework, the partitioned analysis was performed by using a clustering algorithm to group orthologs based on sequence properties and comparing alternative clusters and evolutionary models using the Bayesian Information Criterion (BIC) (Lanfear et al. 2014; Lanfear et al. 2016) (v2.1.1) (--rclusterf), which identified 90 partitions. In two instances, a partition did not contain all 20 amino acid states, which can cause problems with phylogenetic parameter estimation, so for both of these cases, the two partitions were combined with a partition using a similar model of evolution (ortholog 61 combined with ortholog 62 (both JTT), ortholog 71 combined with ortholog 52 (JTT and JTTDCMUT)). We estimated the concatenated ML phylogeny with these partitions and best fitting models of evolution using RAxML (v8.2.12); to ensure a thorough exploration of tree space and support value estimation, we completed 500 bootstrap (BS) replicates with every fifth BS tree used as a starting tree for the ML tree search. To reduce potential effects of saturation and amino acid usage bias (Susko and Roger 2007), which have been shown to exist in crustaceans (Rota-Stabelli, Lartillot, et al. 2013), the concatenated matrix was recoded into Dayhoff6 states (Susko and Roger 2007) (but see Hernandez and Ryan 2021) and a subsequent phylogenetic analysis was carried out with RAxML with a GTR substitution model on the same 88 partitions and an automated bootstrap convergence criterion (autoMRE).

Mixture models were also used for ML and BI tree searches because they have been shown to account for among site variation in amino-acid propensities, and thus are less prone to artifacts like long branch attraction (Lartillot and Philippe 2004; Lartillot et al. 2007; Le et al. 2008), without the information loss inherent in recoding strategies such as Dayhoff6 (Hernandez and Ryan 2021). In the ML mixture model framework, the C60 mixture model LG+C60+F+G was used in IQTREE (v1.6.11) (Nguyen et al. 2015) to build an initial tree, and the resulting tree was used as a guide tree for a posterior mean site frequency model (PMSF) (Wang et al. 2018) (-m LG+C60+F+G -ft) with 100 bootstrap replicates. Additional bootstrap replicates were attempted in the C60 analysis but could not be completed in a reasonable time frame under a complex model (each bootstrap replicate for this dataset took >2 hours to complete on a 40 core compute node). To test for consistency in the C60 analysis, the LG+C60+F+G model was run from a parsimony starting tree and the tree search was repeated five times from random starting trees; all resulting trees were identical, suggesting the ML mixture model tree search was not stuck on a local optimum. To exclude rapidly evolving genes that may exhibit mutational saturation, average branch length was used as a proxy for rate (Oakley et al. 2013) and calculated with a python script that summed branch lengths across each tree and divided by the number of taxa; the 10% (n = 58) fastest evolving genes (those with the highest average branch lengths) were removed from the concatenated ortholog matrix to produce another matrix (LG+C60+F+G – fastest genes).

BI analyses were completed using the CAT-GTR model of PhyloBayes-MPI (v1.8) (Lartillot et al. 2013) with three independent chains. Each chain was run for over 35,000 generations (>6 months of continual run time on 2×40 core compute nodes for >300,000 CPU hours of runtime for each chain). Convergence between the three chains in the BI analysis were assessed using the PhyloBayes bpcomp module sampling every 10 trees with the first 25% of trees excluded as burn-in. Support values were obtained by calculating the posterior probability at each node.

For the multispecies coalescent phylogeny, individual gene trees were built for each ortholog using IQTREE with the amino acid substitution model of best fit by BIC score and 200 BS replicates. The species tree was estimated by using the ML gene trees as input in ASTRAL-III (v5.6.3) (Mirarab et al. 2016; Zhang et al. 2018). The sensitivity of the ASTRAL-III analyses to a number of ortholog features was explored. Ortholog features were based on Shen et al. (2016), which identified gene properties most strongly associated with phylogenetic signals. These properties were measured using the python package ETE3 (Huerta-Cepas et al. 2016) unless otherwise noted. The following were measured for each ortholog: number of taxa, alignment length, total amino acids in alignment, number of gaps in alignment, percent of gaps, number of variable sites, proportion of variable sites, number of parsimony informative sites, proportion of parsimony informative sites, tree length, average branch length (tree length / number of taxa), and compositional homogeneity (Table S4). ML ortholog branches with less than 10% and 30% BS support in the individual ortholog trees were also collapsed prior to running ASTRAL-III following Zhang et al. (2018). Branch support for the ASTRAL-III analyses was assessed using local posterior probabilities (Sayyari and Mirarab 2016).

#### Comparison of phylogenetic results

We identified shared orthogroups between the datasets using BLASTp. Specifically, a reciprocal BLASTp analysis was performed between datasets with the amino acid sequences that had not been trimmed for the phylogenetic analyses to avoid missing amino acid sites. Because we predicted orthologs using maximum inclusion and the different datasets include different taxa, we only considered orthogroups that share the same sequences (i.e., 100% identity) and the same taxa. The different taxa in each dataset ultimately affect the MCL homolog clustering and ortholog prediction; therefore, we only chose to compare orthologs from different datasets with the same taxa and sequences.

Topological differences from the phylogenetic analyses were assessed using Robinson-Foulds (RF) symmetric distances calculated in ETE3 (Huerta-Cepas et al. 2016) and the information metric of Kendall and Colijn (2016) using the R package TreeSpace (Jombart et al. 2017). A multidimensional scaling (MDS) plot showing topological variation between analyses based on the Kendall and Colijn metric was also calculated with TreeSpace (Fig. 3). Topologies from the different phylogenetic analyses were compared by AU-test (Shimodaira 2002) in IQTREE (v1.6.11) along with published pancrustacean phylogenies (Regier et al. 2010; von Reumont et al. 2012; Oakley et al. 2013; Rota-Stabelli, Lartillot, et al. 2013; Schwentner et al. 2017; Schwentner et al. 2018; Lozano-Fernandez et al. 2019) using the full ortholog matrix and an LG+G+F+I model with 100,000 BS with the RELL method.

We also estimated Long Branch (LB) scores with Phykit v1.11.10 (Steenwyk et al. 2021) to compare the relative lengths of terminal branches in each phylogeny. The LB score for each taxon is the mean patristic distance between itself and all other taxa directly proportional to the mean of all patristic distances for all taxa in the phylogeny (Struck 2014; Weigert et al. 2014). The larger the LB score, the longer the terminal branch with respect to all other taxa in the tree. While this metric serves as a measure to compare terminal branches in a phylogeny, it cannot be used here to compare terminal branches in different phylogenies because branch lengths are parameter estimates and our model changed in each phylogeny due to different taxon and gene sampling.

### Divergence time estimation

Divergence time estimation was based on 12 vetted internal fossil calibrations (six from Wolfe et al. (2016), two from Wolfe et al. (2019), and four new) and the root prior was defined based on the Euarthropoda node (Wolfe et al. 2016, node 4) with a gamma distribution with mean 575 Ma and SD 61 Ma (fossil ages and justifications in Supplemental Table and text S7). Divergence times for the main analysis were estimated using MCMCTree (dos Reis and Yang 2011; dos Reis et al. 2015) and the full aa matrix of Dataset 3 using the fixed topology of the highest likelihood tree, which resulted from the LG+C60+F+G analysis. Divergence times were calculated using an independent, log-normal model (clock = 2) and approximate likelihood calculation; the Hessian calculation for approximate likelihood was done using an LG+G4 matrix (LG was the model of best fit by AIC and BIC). We ran 3 chains for 20 million generations each, treating the first 25% as burnin and sampling every 1,000 trees. A fourth chain with the same settings without data to sample from the time prior. Convergence of the 3 chains was assessed visually by plotting the distributions of the 3 chains against each other. The distributions of the chains were nearly perfectly linear at 2 million generations. To further ensure convergence, we ran each chain for 20 million generations and assessed convergence visually in the same way (Figure S7). Results in Figures 4 and S9 were plotted using the *MCMCtreeR* package (Puttick 2019).

Divergence times were also estimated in PhyloBayes v1.8 (Lartillot et al. 2013) using a fixed topology from the C60+LG+F+G analysis. Due to the size of our data matrices and time to convergence, we assembled an amino acid alignment consisting of the 50 loci with the highest normalized Robinson-Foulds distance compared to the species tree resulting from the LG+C60+F+G analysis of the full matrix. This subsampled matrix was then used for divergence time estimation in PhyloBayes (Mongiardino Koch 2021). The inability to use the entire alignment is why we included these as supplementary analyses. We used the C20+LG substitution model and compared the uncorrelated gamma multipliers (UGM) and lognormal (LN) relaxed clock models (Drummond et al. 2006) and the autocorrelated CIR clock model (Lepage et al. 2007) with 3 chains per run. Although the topology was fixed, we used a birth-death tree model, with soft bounds allowing 5% of the probability distribution outside the input fossil ages. An automatic stopping rule was implemented, with tests of convergence every 100 cycles, until the default criteria of effective sample sizes and parameter discrepancies between chains were met (50 and 0.3, respectively). While many values did converge to <0.3, as is commonly the case with PhyloBayes, not all values fully converged even after months of runtime (sigma, mu, scale, and p2 were consistently >0.3 while all other parameters were <0.1). To further assess convergence, we visualized trace plots of logL values for each of the 3 chains running for each model in R (Table S8). Trees and their posterior distributions were generated from completed chains after the initial 20% of sampled generations were discarded. We compared estimated posterior age distributions to the marginal prior by removing sequence data using the -prior flag (Warnock et al. 2012; Brown and Smith 2017).

## Data availability

All alignments and tree files from this study are available on Dryad: https://doi.org/10.5061/dryad.dr7sqvb2h

## Acknowledgements

We thank Nicolás Mongiardino Koch for helpful discussions on divergence time estimation and for a script for generating logL plots of PhyloBayes chains. We are grateful to David Fenwick and Aphotomarine.com for the use of their buoy barnacle photograph. This material is based in part upon work supported by the NSF PRFB Program #2010898 to JPB. JMW is supported by NSF DEB #1856679.

## Supplemental Information

**Table S1**. Genome and transcriptome sample information and ortholog occupancy for Dataset 3.

**Table S2**. Genome and transcriptome sample information and ortholog occupancy for Dataset 2.

**Table S3**. Genome and transcriptome sample information and ortholog occupancy for Dataset 1.

**Table S4**. Ortholog properties.

**Table S5**. Long Branch scores for taxa in Datasets 2 and 3.

**Table S6**. Pancrustacean topology tests.

**Table S7**. Fossil dates table and justifications.

**Figure S1.**
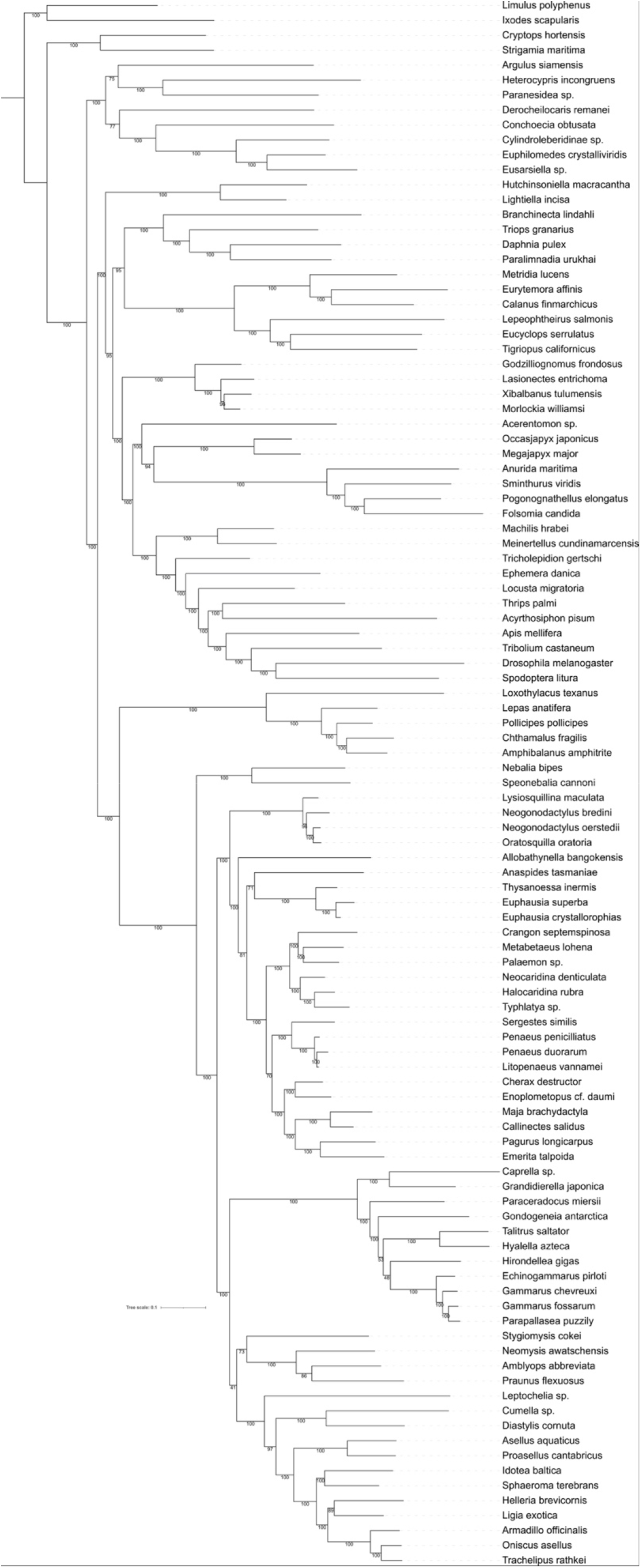
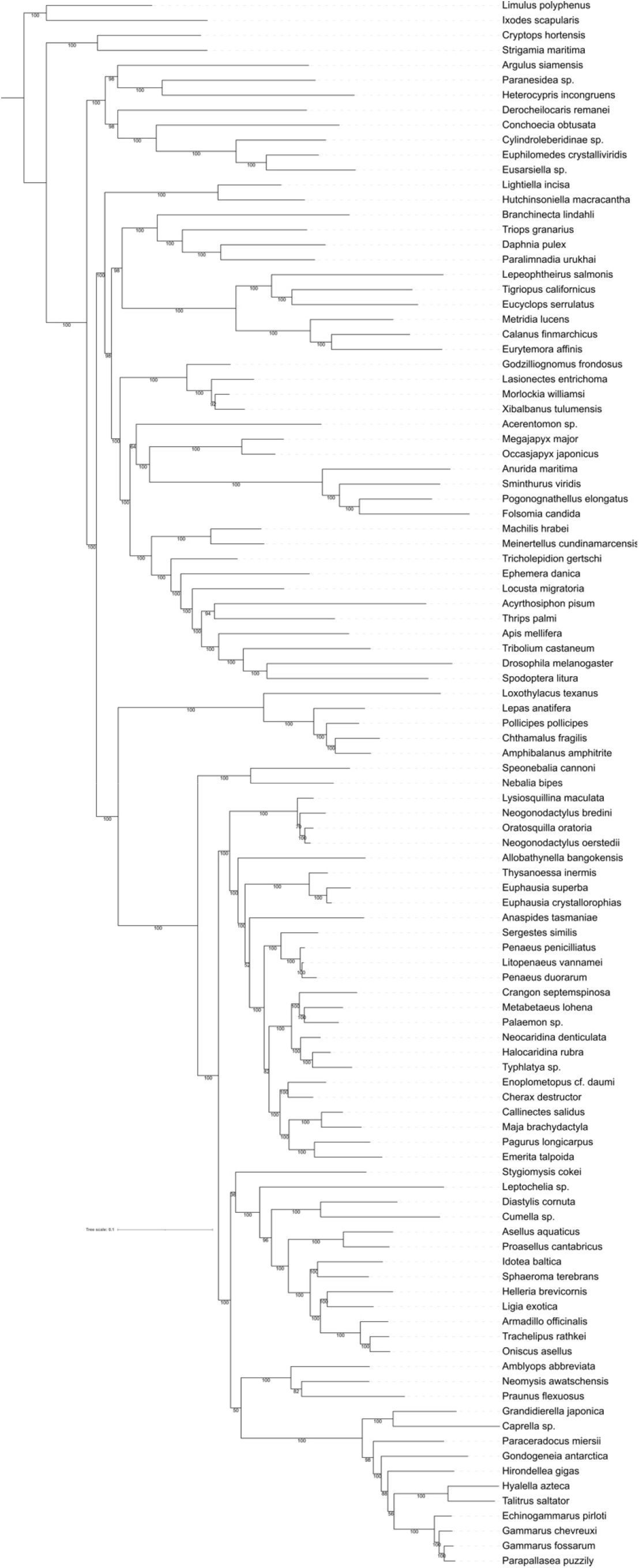
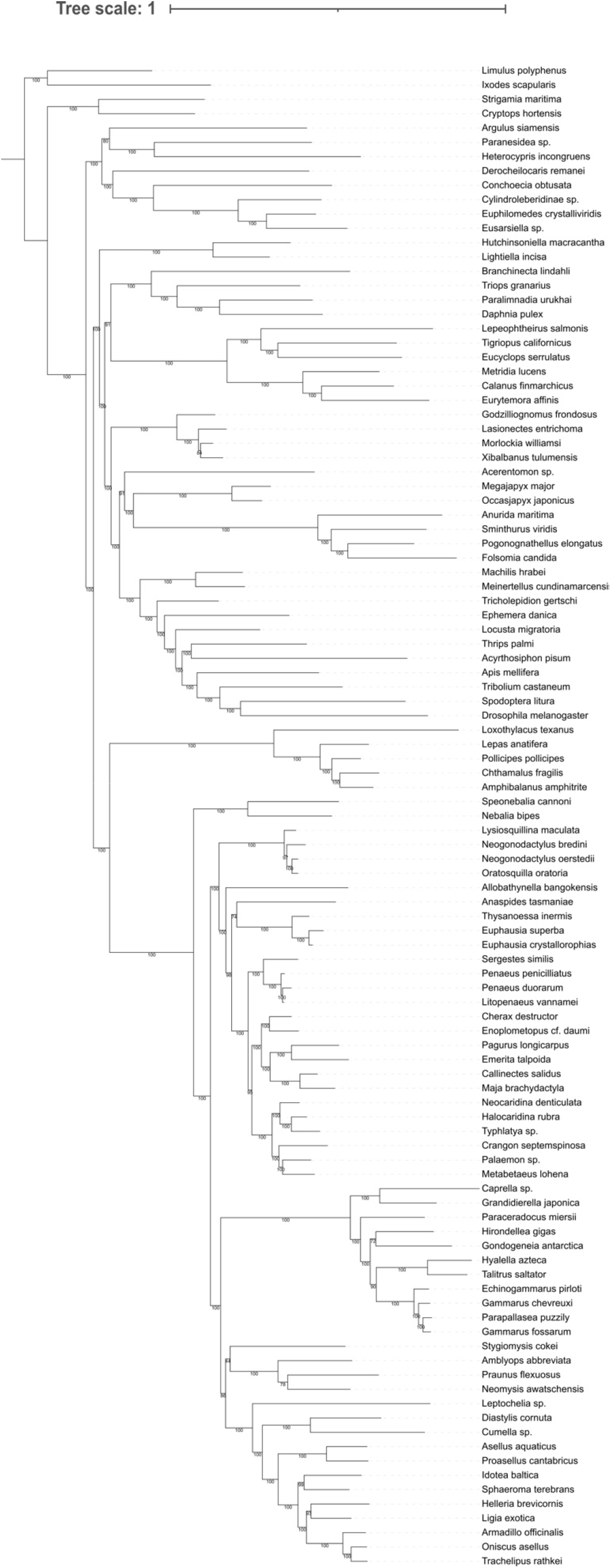
Other ML topologies (A) RAxML partitioned analysis; (B) RAxML Dayhoff6 recoded matrix; (C) LG+C60+F+G with 10% fastest evolving genes removed from matrix.

**Figure S2.**
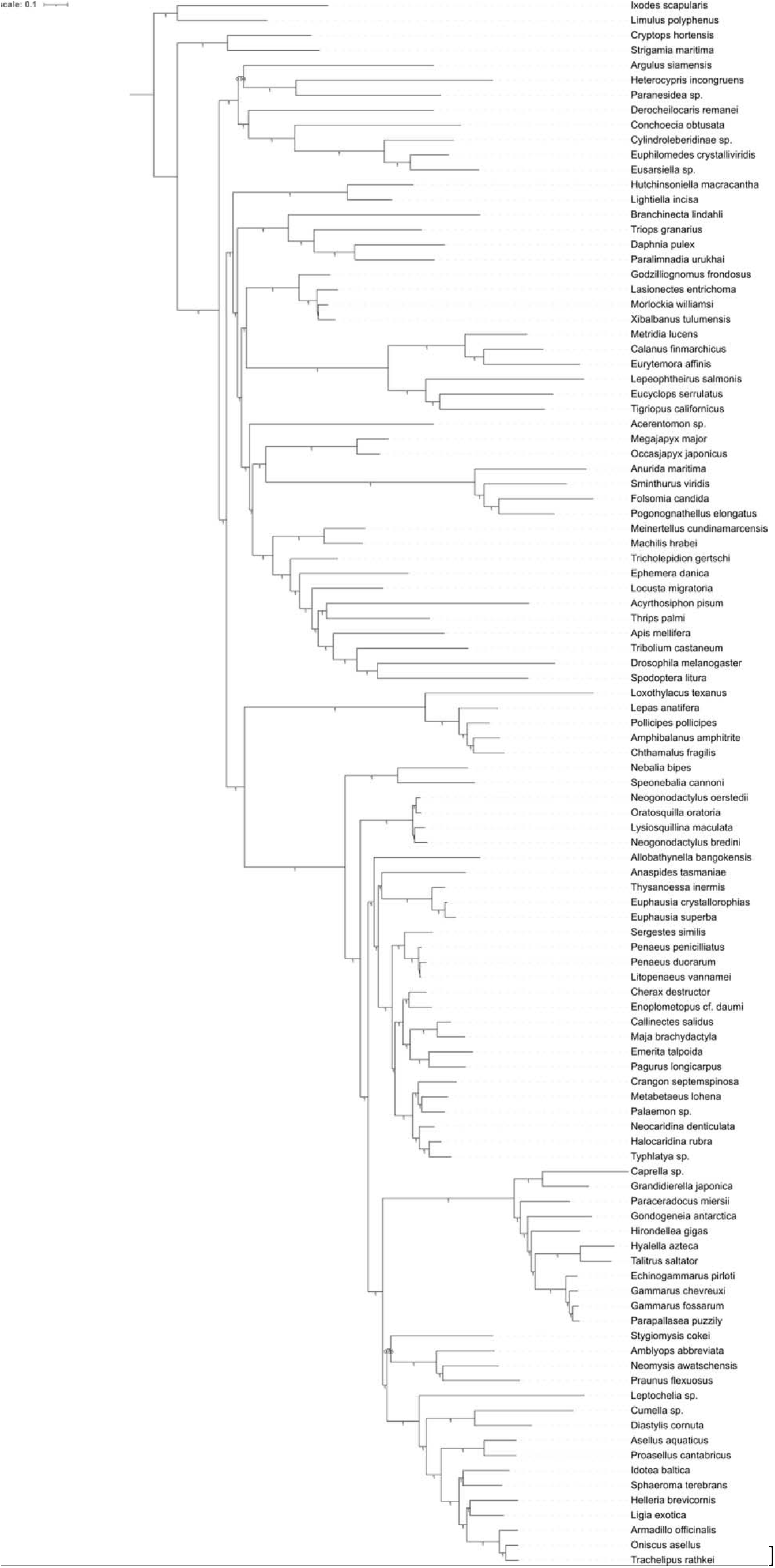
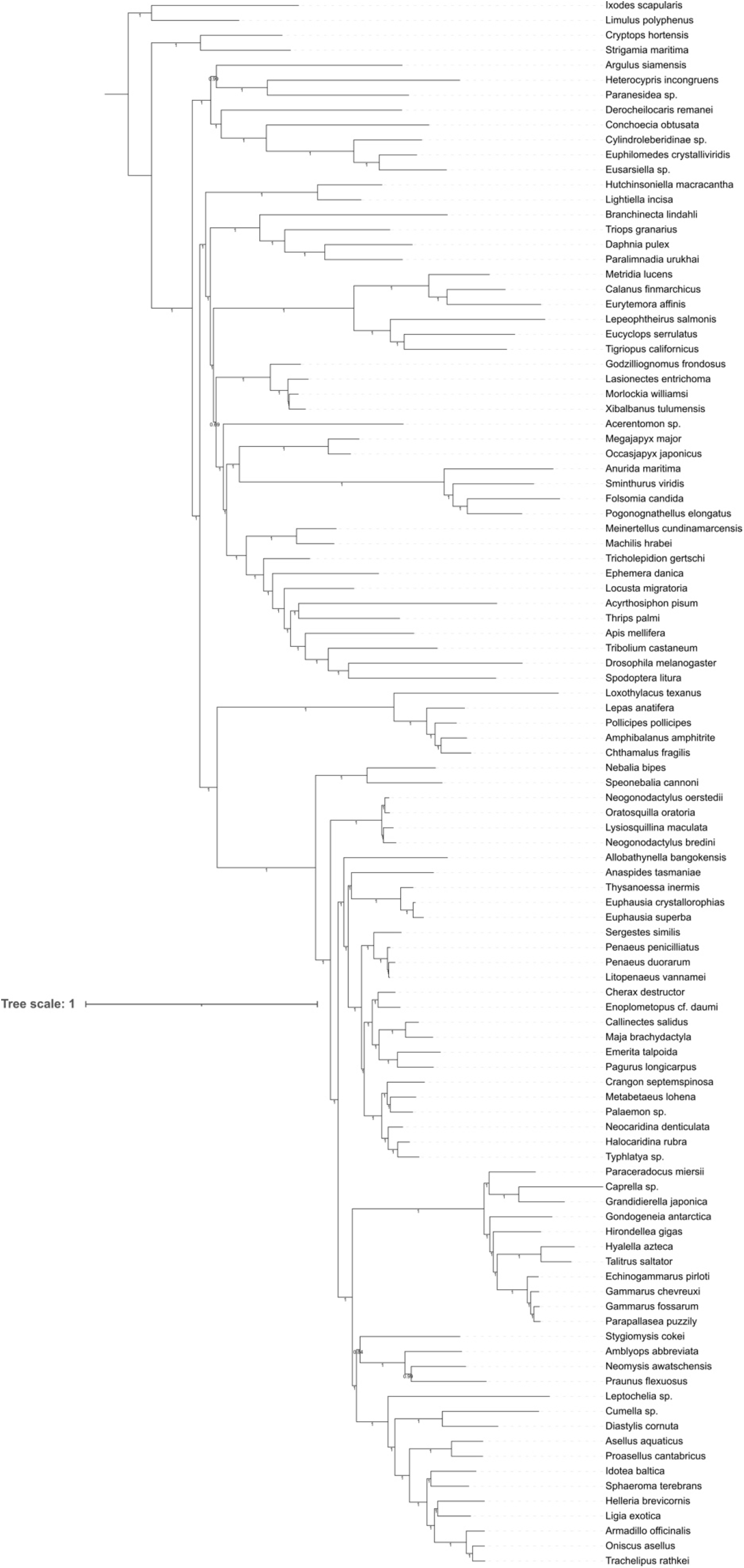
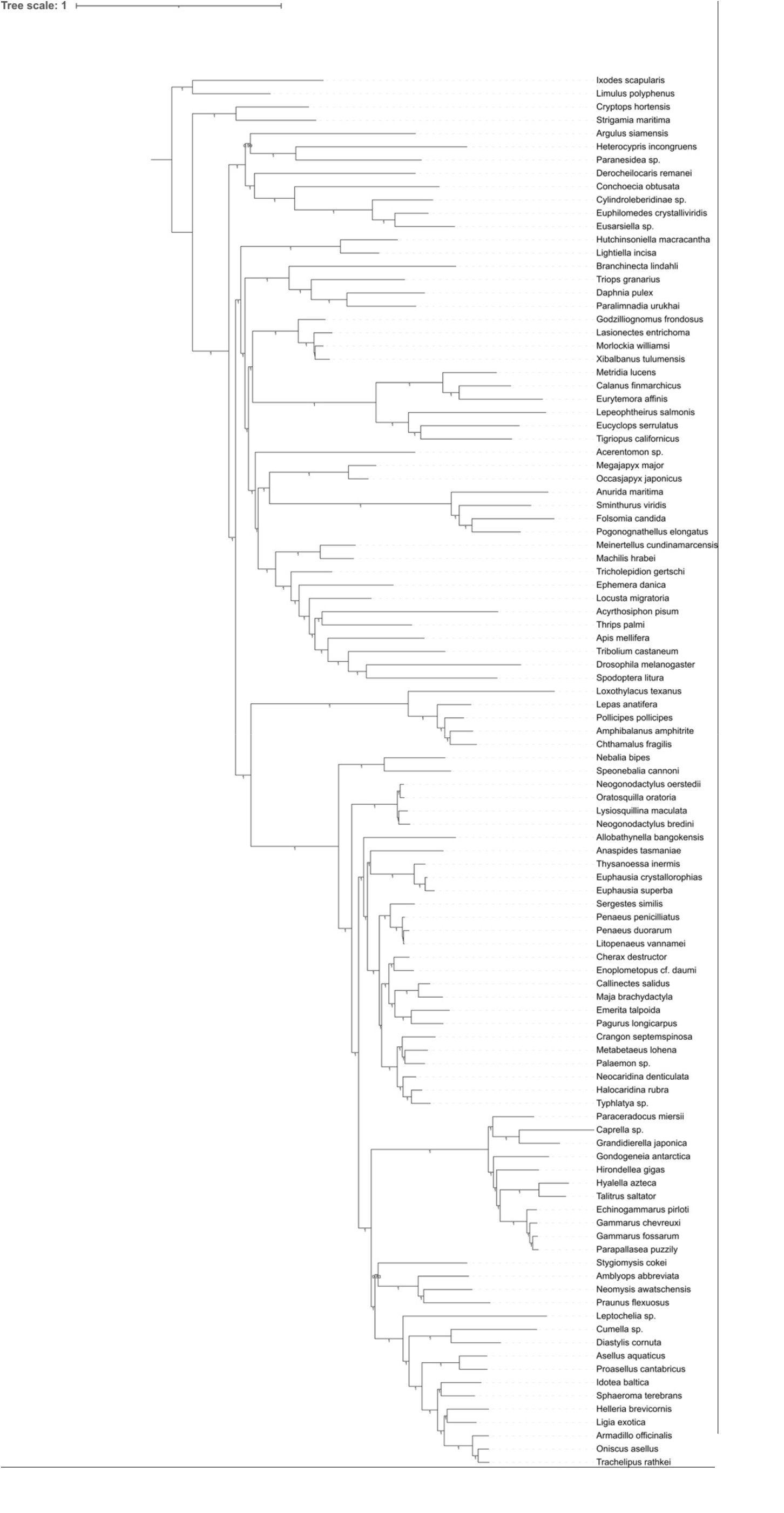
**Phylobayes** (A) chain 1; (B) chain 2; (C) chain 3.

**Figure S3.**
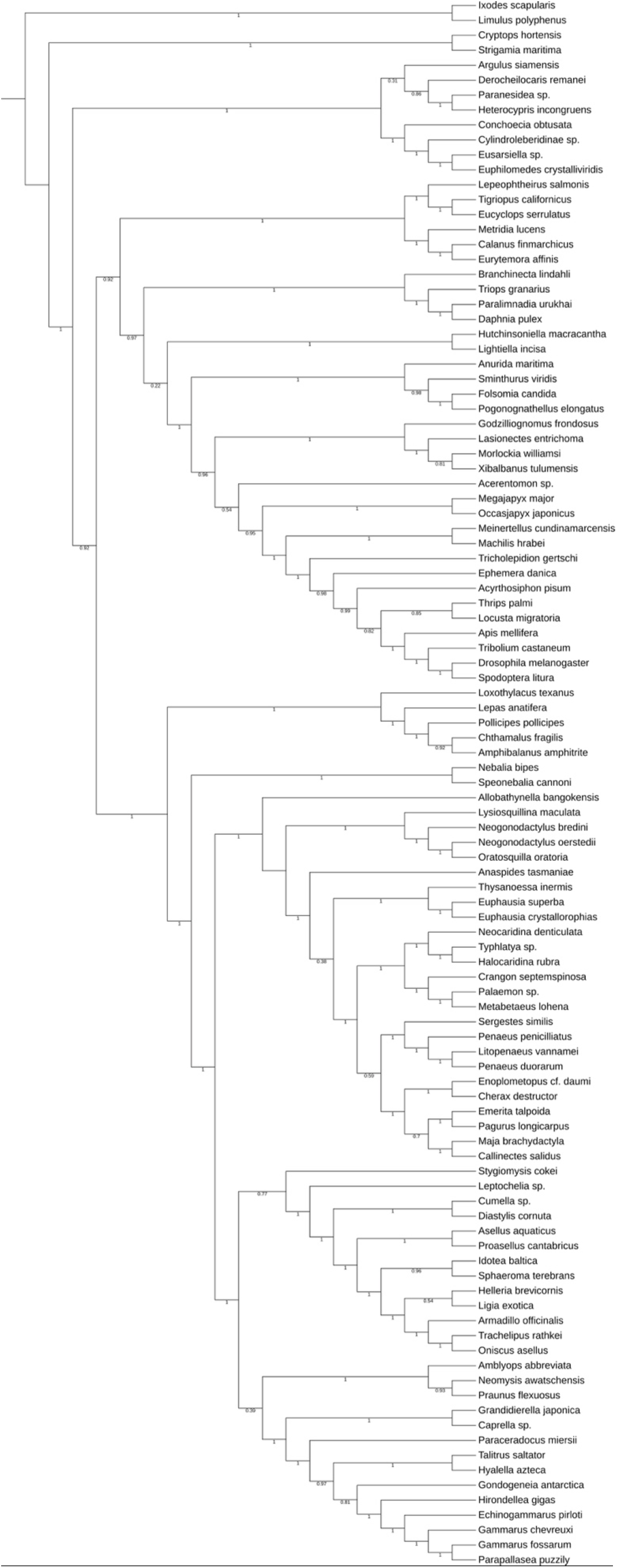
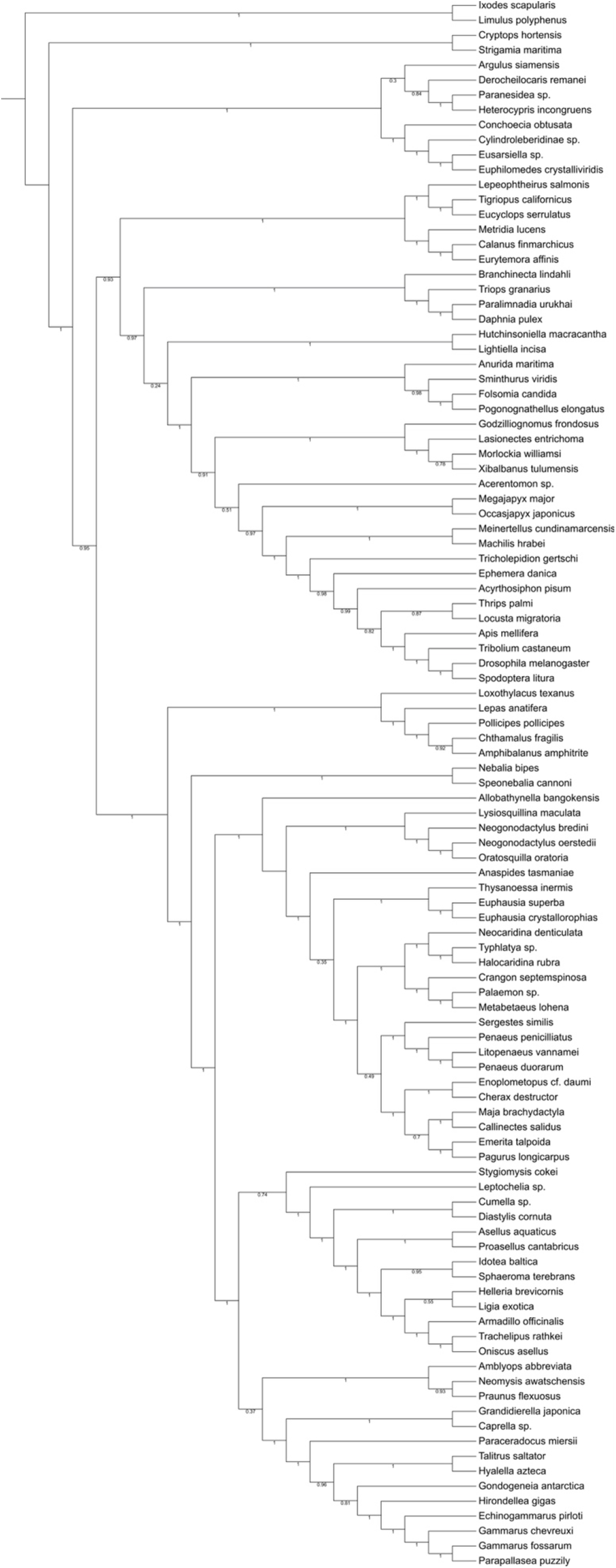
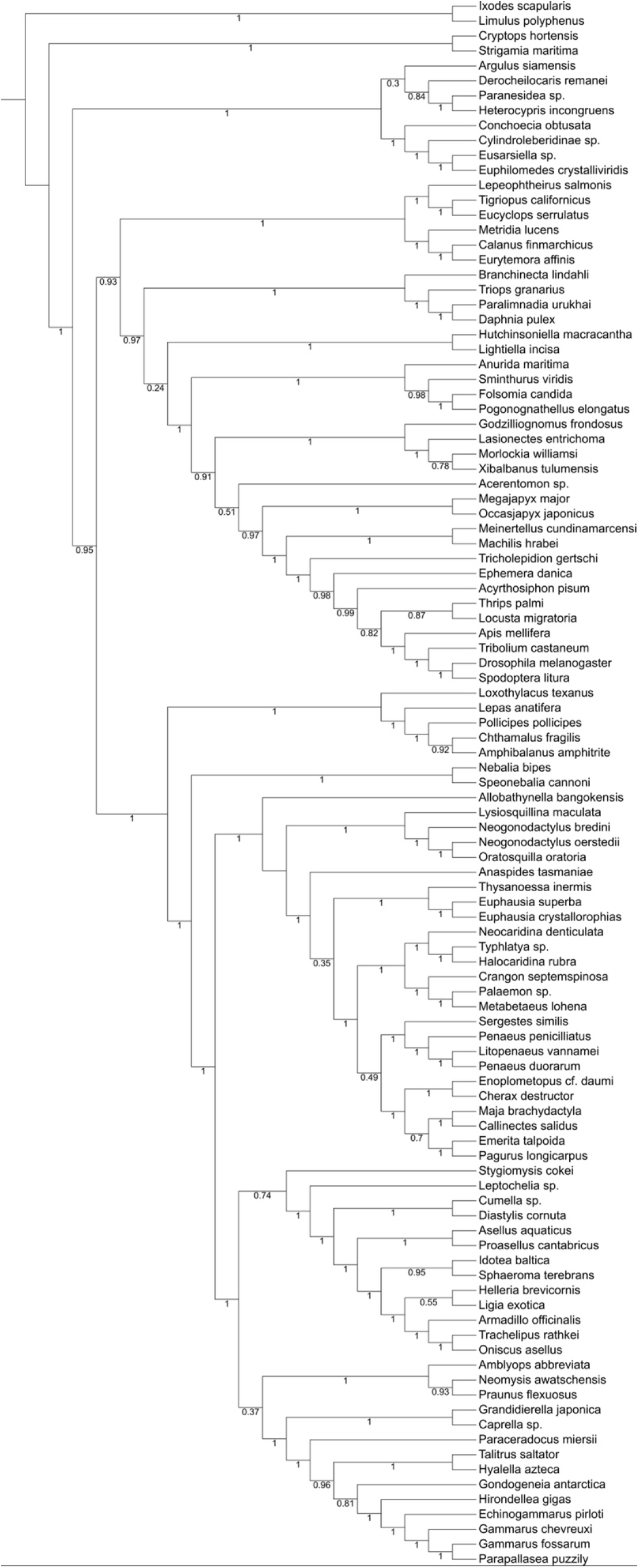
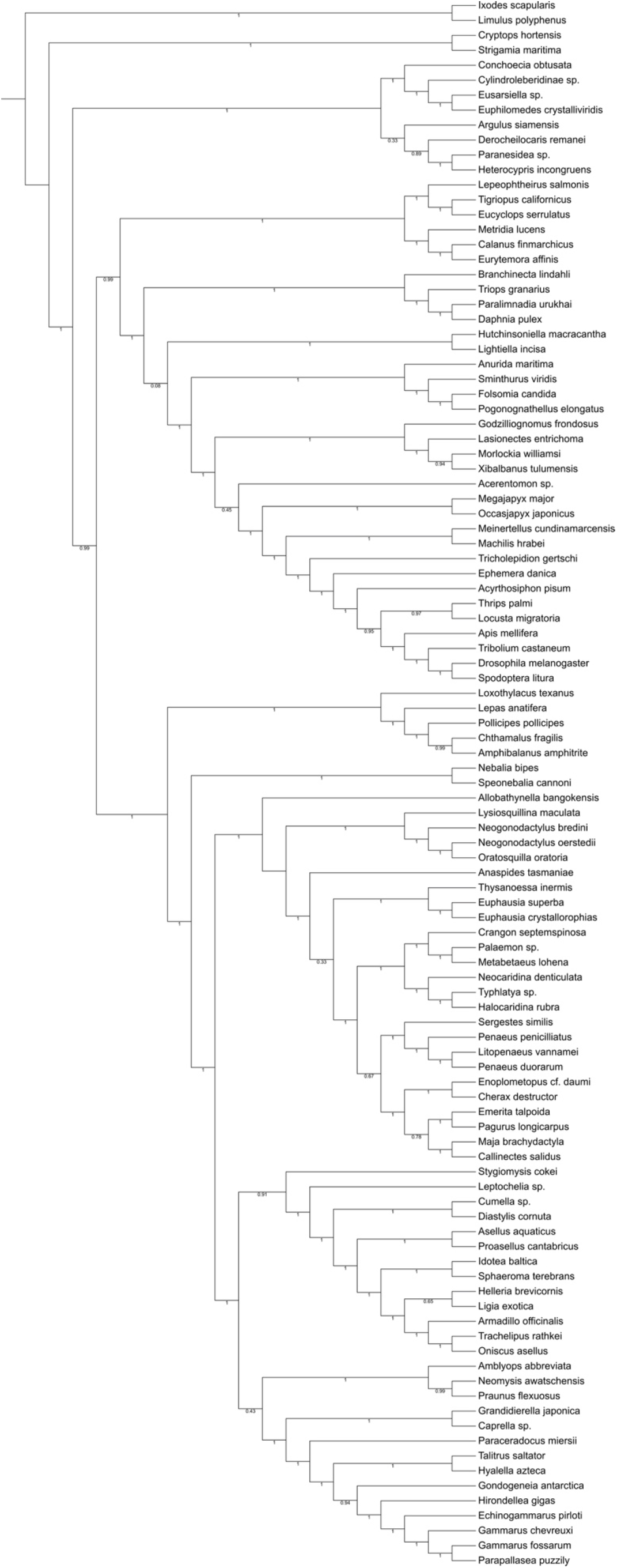
**ASTRAL** (A) all nodes in gene trees; (B) gene tree nodes <10% BS support collapsed; (C) gene tree nodes <30% BS support collapsed.

**Figure S4.**
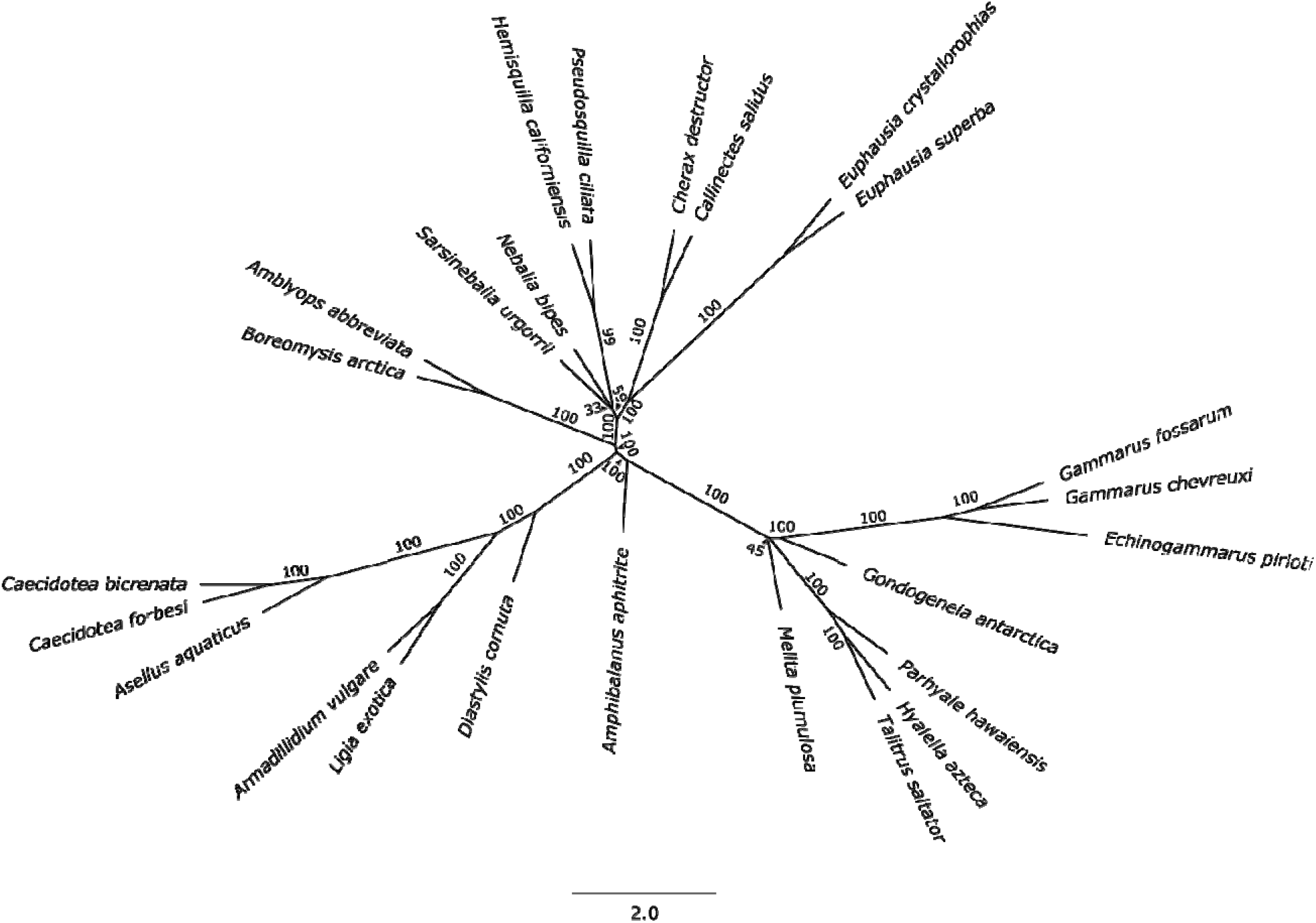
Tree results from ASTRAL analyses of Dataset 1.

**Figure S5.**
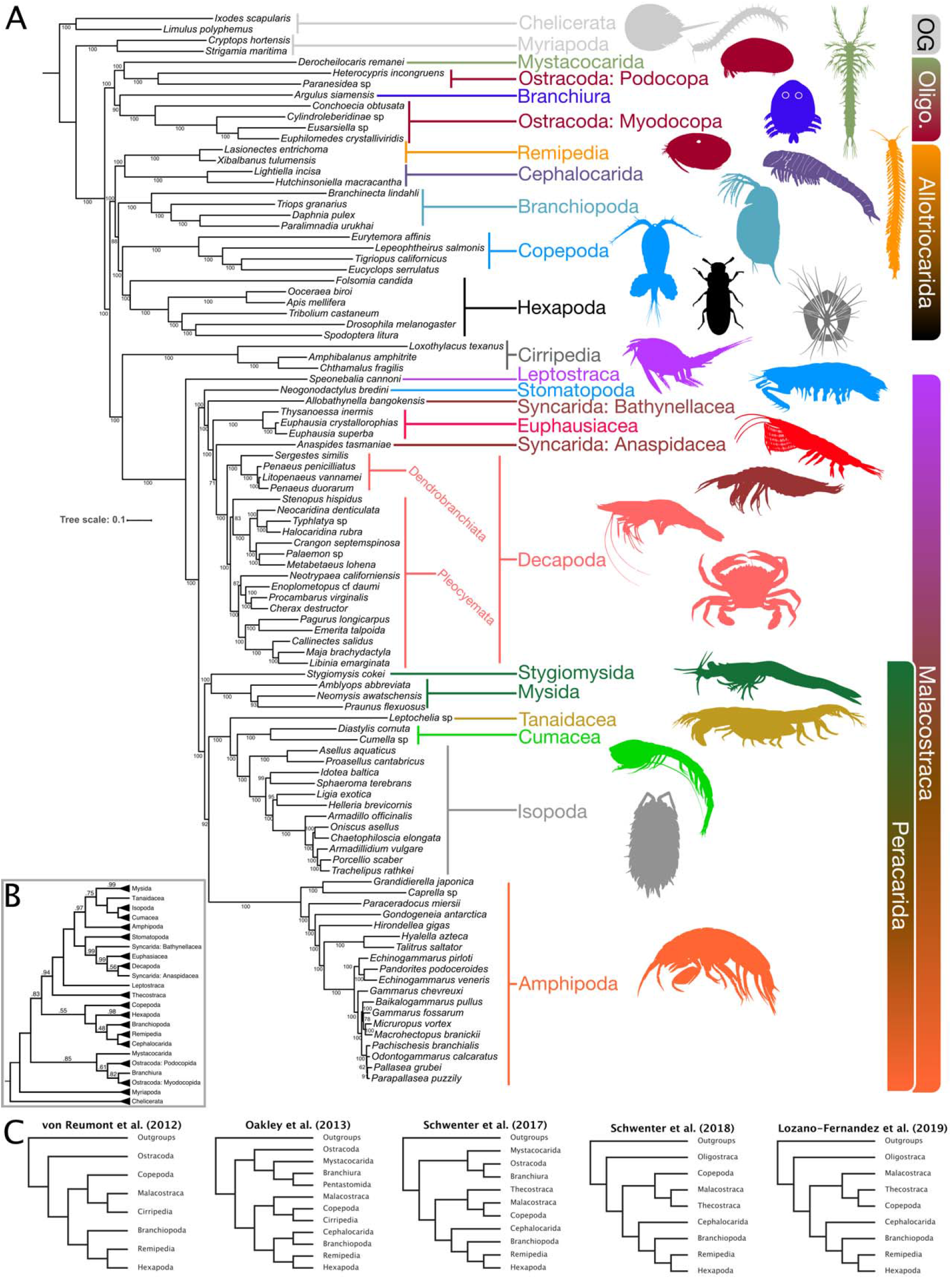
Tree resulting from LG+C60+F+G analyses of Dataset 2.

**Figure S6.**
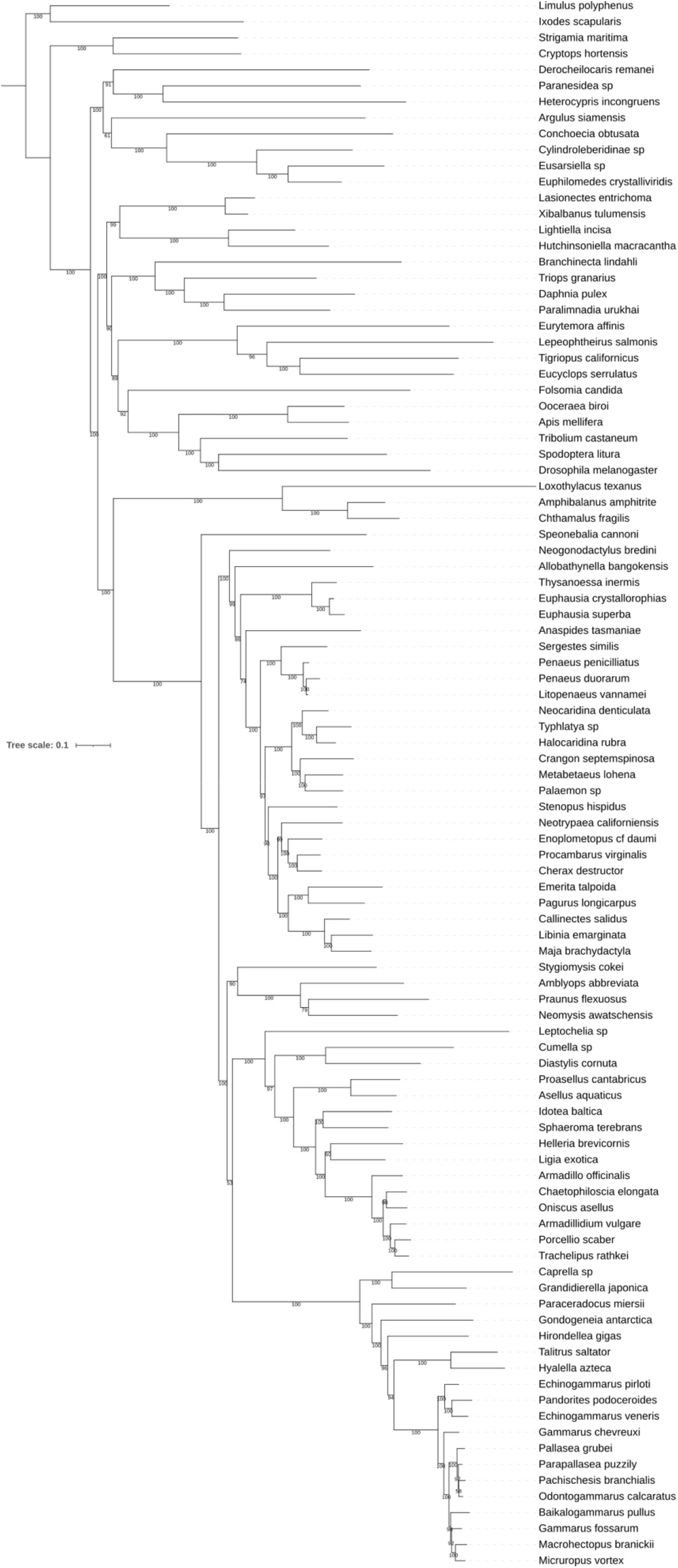
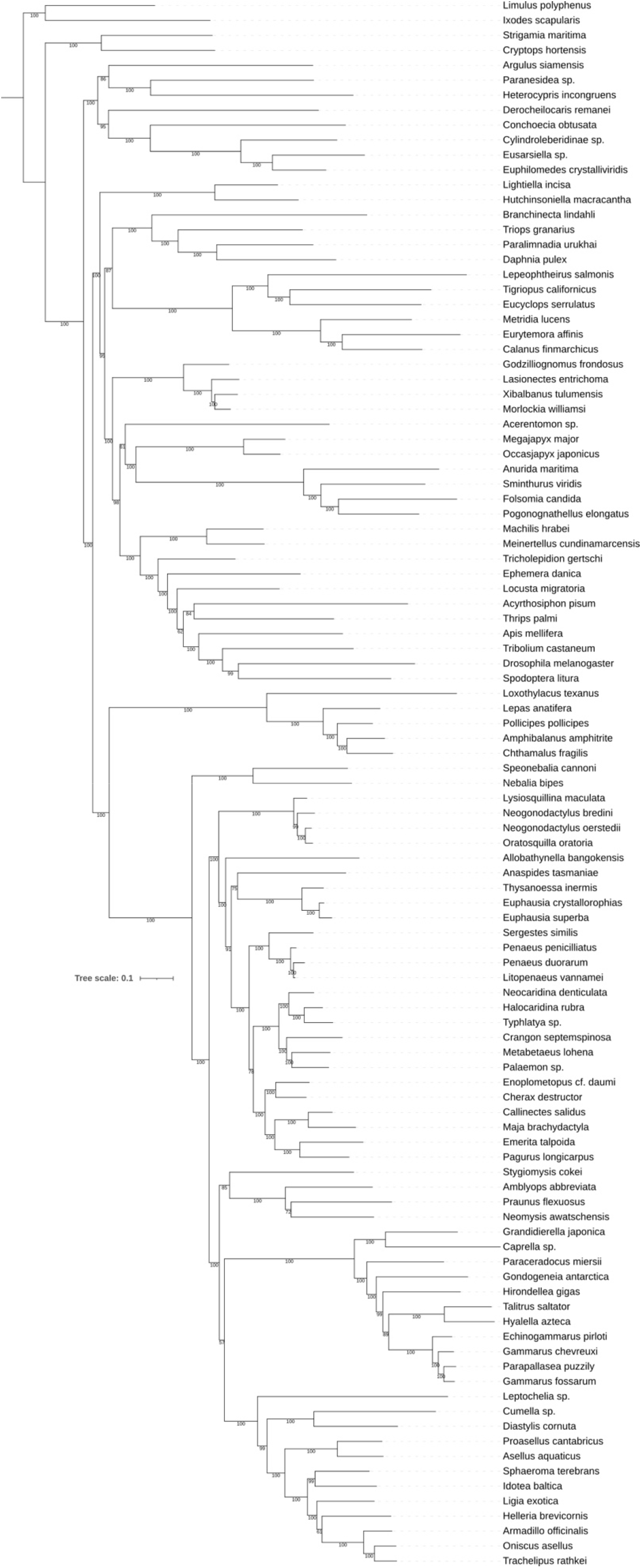
Trees results from LG+C60+F+G analysis taxa from Datasets 2 and 3, using the same orthologs (only genes shared between the datasets). (A) Dataset 2 taxa; (B) Dataset 3 taxa.

**Figure S7.**
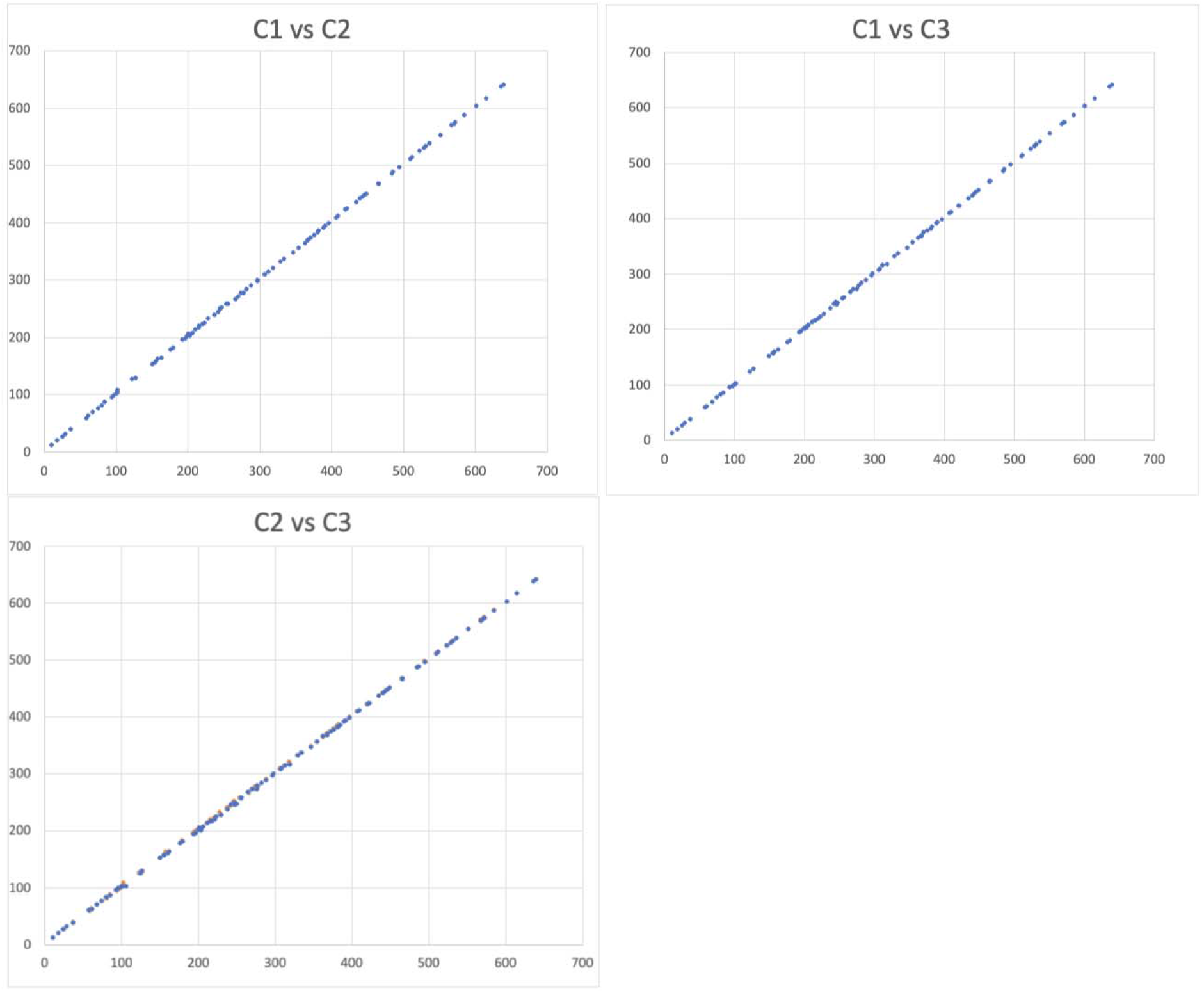
Assessment of convergence of divergence time estimation analysis of 3 MCMCTree chains by plotting posterior mean times from 20 million generations.

**Figure S8.**
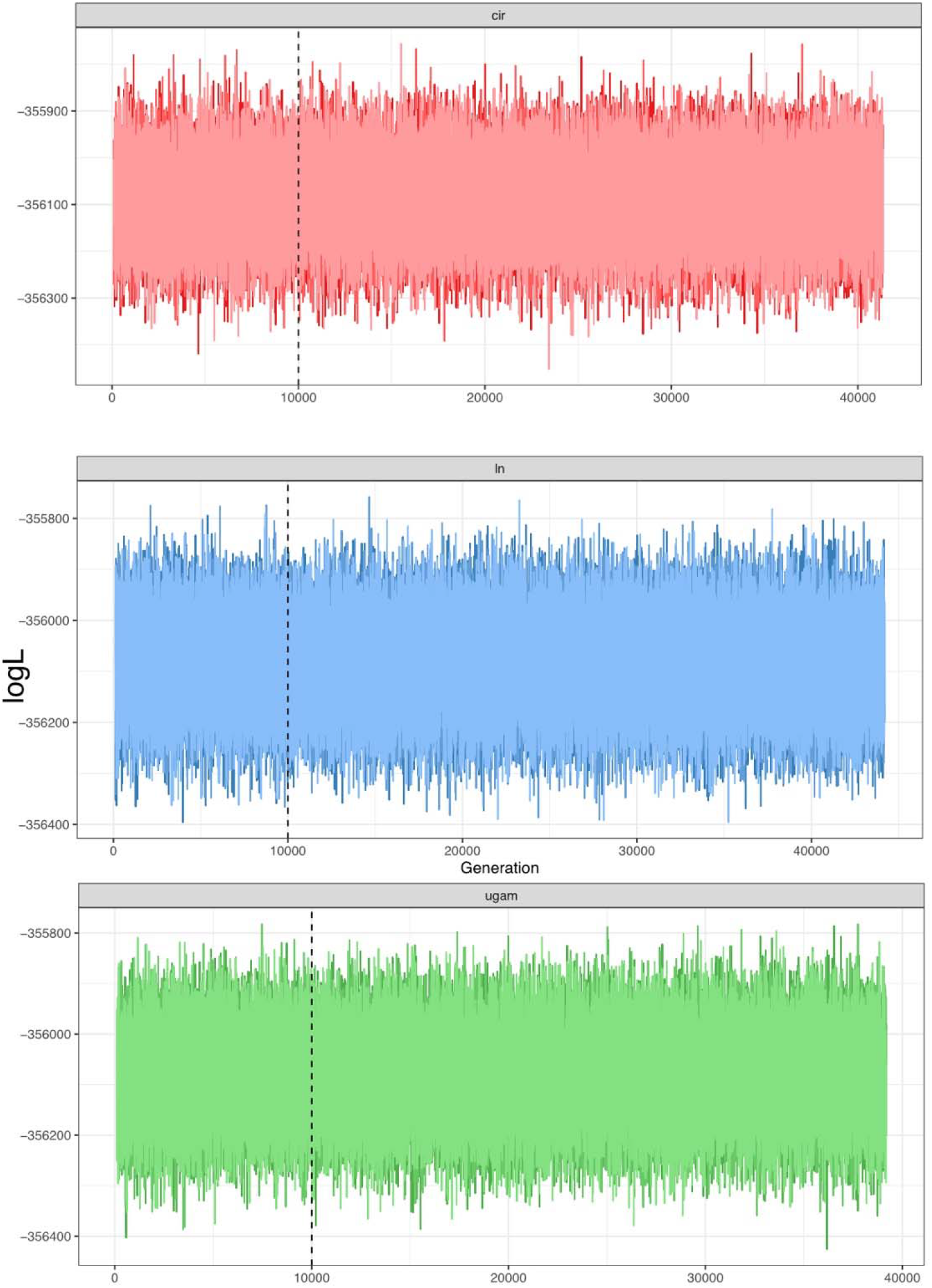
Traceplots of 3 Phylobayes chains each for CIR, LN, and UGAM divergence time analses after approximately 40,000 generations.

**Figure S9.**
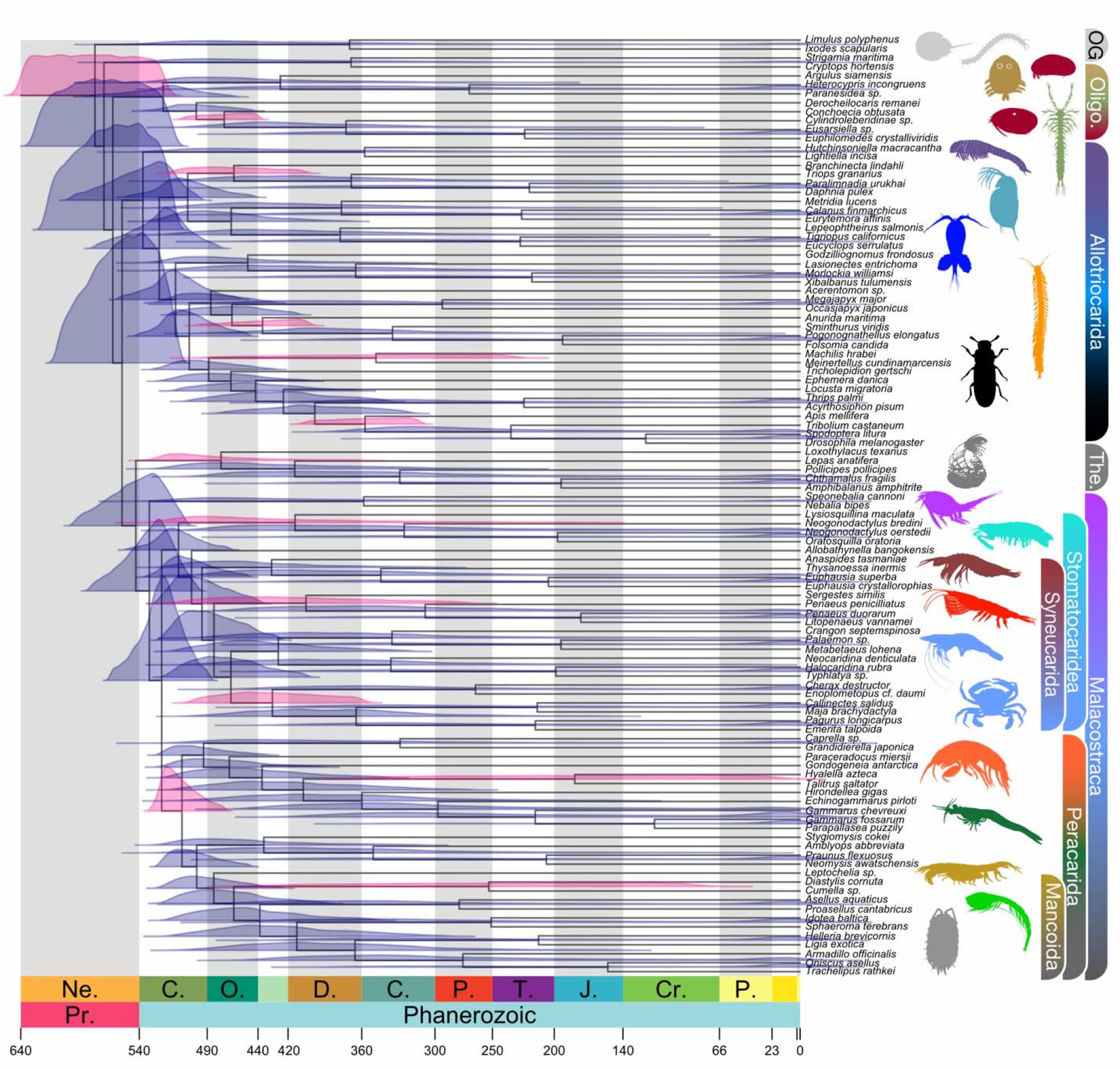
Fossil calibrated divergence time estimates under the marginal prior (no sequence data), based on an MCMCtree analysis of the topology depicted in Figure 2A calibrated with 13 vetted fossils.

**Figure S10.**
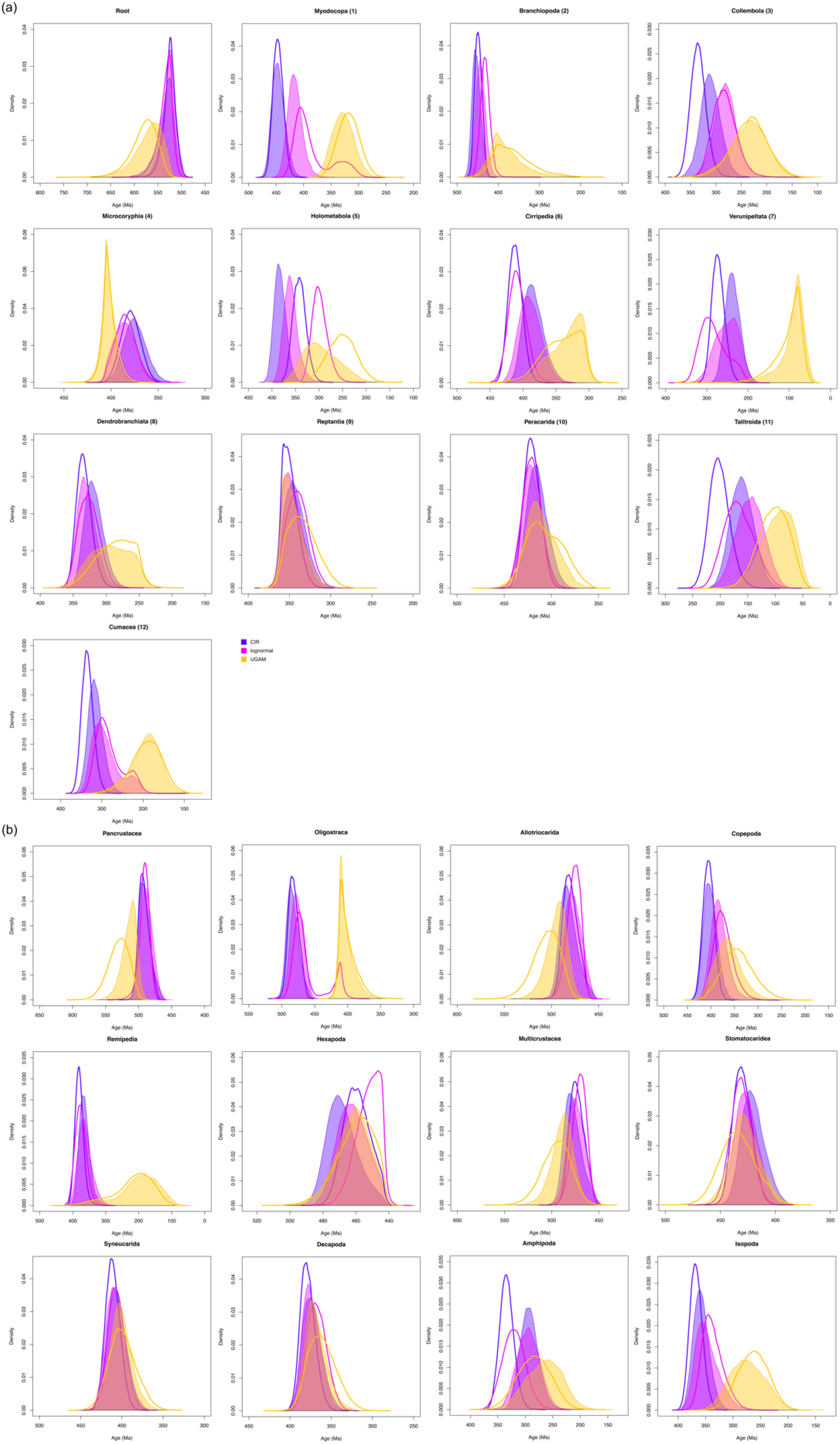
Comparison of posterior probability distributions for divergence time analyses under different clock models in PhyloBayes, and the same analyses under the marginal prior (removing sequence data). The posterior analyses are shaded; marginal priors are superimposed on the same axes with a heavy line of the same color. (A) All nodes directly calibrated by fossils and their calibration number; (B) Selected nodes calibrated by only the tree prior.

## References

Alfsnes K, Leinaas HP, Hessen DO. 2017. Genome size in arthropods; different roles of phylogeny, habitat and life history in insects and crustaceans. Ecology and Evolution 7:5939–5947.

Altenhoff AM, Dessimoz C. 2009. Phylogenetic and Functional Assessment of Orthologs Inference Projects and Methods. PLoS Comput Biol 5:e1000262.

Altenhoff AM, Schneider A, Gonnet GH, Dessimoz C. 2011. OMA 2011: orthology inference among 1000 complete genomes. Nucleic Acids Research 39:D289–D294.

Altschul S, Gish W, Miller W, Myers EW, Lipman DJ. 1990. Basic Local Alignment Search Tool. Journal of Molecular Biology 215:403–410.

Altschul S, Madden TL, Schäffer AA, Zhang J, Zhang Z, Miller W, Lipman DJ. 1997. Gapped BLAST and PSI-BLAST: a new generation of protein database search programs. Nucleic Acids Research 25:3389–3402.

Andrew DR. 2011. A new view of insect–crustacean relationships II. Inferences from expressed sequence tags and comparisons with neural cladistics. Arthropod Structure & Development 40:289–302.

Andrews S. 2018. FastQC: a quality control tool for high throughput sequence data. Available from: http://www.bioinformatics.babraham.ac.uk/projects/fastqc/

Ballesteros JA, Hormiga G. 2016. A new orthology assessment method for phylogenomic data: Unrooted Phylogenetic Orthology. Molecular Biology and Evolution 33:2117–2134.

Barba-Montoya J, dos Reis M, Yang Z. 2017. Comparison of different strategies for using fossil calibrations to generate the time prior in Bayesian molecular clock dating. Molecular Phylogenetics and Evolution 114:386–400.

Bar-On YM, Phillips R, Milo R. 2018. The biomass distribution on Earth. Proceedings of the National Academy of Sciences 115:6506–6511.

Bayzid MS, Warnow T. 2013. Naive binning improves phylogenomic analyses. Bioinformatics 29:2277–2284.

Bernot JP, Avdeyev P, Zamyatin A, Dreyer N, Alexeev N, Pérez-Losada M, Crandall KA. 2022. Chromosome-level genome assembly, annotation, and phylogenomics of the gooseneck barnacle Pollicipes pollicipes. GigaScience 11:giac021.

Betancur-R. R, Arcila D, Vari RP, Hughes LC, Oliveira C, Sabaj MH, Ortí G. 2019. Phylogenomic incongruence, hypothesis testing, and taxonomic sampling: The monophyly of characiform fishes. Evolution 73:329–345.

Bolger AM, Lohse M, Usadel B. 2014. Trimmomatic: a flexible trimmer for Illumina sequence data. Bioinformatics 30:2114–2120.

Bracken-Grissom H, Wolfe JM. 2020. The pancrustacean conundrum: a conflicted phylogeny with emphasis on Crustacea. In: Poore GCB, Thiel M, editors. Evolution and Biogeography. Vol. 8. Natural History of the Crustacea. Oxford University Press. p. 28.

Branstetter MG, Danforth BN, Pitts JP, Faircloth BC, Ward PS, Buffington ML, Gates MW, Kula RR, Brady SG. 2017. Phylogenomic Insights into the Evolution of Stinging Wasps and the Origins of Ants and Bees. Current Biology 27:1019–1025.

Brown J, Smith S. 2017. The Past Sure Is Tense: On Interpreting Phylogenetic Divergence Time Estimates. Systematic Biology 67:340–353.

Budd GE, Mann RP. 2020a. The dynamics of stem and crown groups. Science Advances 6:eaaz1626.

Budd GE, Mann RP. 2020b. Survival and selection biases in early animal evolution and a source of systematic overestimation in molecular clocks. Interface Focus 10:20190110.

Camacho C, Coulouris G, Avagyan V, Ma N, Papadopoulos J, Bealer K, Madden TL. 2009. BLAST+: architecture and applications. BMC Bioinformatics 10:421.

Chan, BK, Dreyer, N, Gale, AS, Glenner, H, Ewers-Saucedo, C, Pérez-Losada, M, Kolbasov, GA, Crandall, KA and Høeg, JT, 2021. The evolutionary diversity of barnacles, with an updated classification of fossil and living forms. Zoological Journal of the Linnean Society, 193(3): 789–846.

Chen F, Mackey AJ, Vermunt JK, Roos DS. 2007. Assessing Performance of Orthology Detection Strategies Applied to Eukaryotic Genomes. PLoS ONE 2:e383.

Chiu JC, Lee EK, Egan MG, Sarkar IN, Coruzzi GM, DeSalle R. 2006. OrthologID: Automation of genome-scale ortholog identification within a parsimony framework. Bioinformatics 22:699–707.

Daley AC, Antcliffe JB, Drage HB, Pates S. 2018. Early fossil record of Euarthropoda and the Cambrian Explosion. Proceedings of the National Academy of Sciences of the United States of America 115:5323–5331.

DeGiorgio M, Degnan JH. 2014. Robustness to Divergence Time Underestimation When Inferring Species Trees from Estimated Gene Trees. Systematic Biology 63:66–82.

Drummond AJ, Ho SYW, Phillips MJ, Rambaut A. 2006. Relaxed Phylogenetics and Dating with Confidence. PLoS Biology 4:e88.

Dunn CW, Hejnol A, Matus DQ, Pang K, Browne WE, Smith SA, Seaver E, Rouse GW, Obst M, Edgecombe GD, et al. 2008. Broad phylogenomic sampling improves resolution of the animal tree of life. Nature 452:745–749.

Dunn CW, Howison M, Zapata F. 2013. Agalma: an automated phylogenomics workflow. BMC Bioinformatics 14:330.

Ellis, EA, Goodheart, JA, Hensley, NM, González, VL, Reda, NJ, Rivers, TJ, Morin, JG, Torres, E, Gerrish, GA, Oakley, TH. 2022. Sexual signals persist over deep time: ancient co-option of bioluminescence for courtship displays in cypridinid ostracods. Systematic Biology: syac057,

Fishbein M, Hibsch-Jetter C, Soltis DE, Hufford L. 2001. Phylogeny of Saxifragales (Angiosperms, Eudicots): Analysis of a Rapid, Ancient Radiation. Systematic Biology 50:817–847.

Fu L, Niu B, Zhu Z, Wu S, Li W. 2012. CD-HIT: accelerated for clustering the next-generation sequencing data. Bioinformatics 28:3150–3152.

Gelman A, Rubin DB. 1992. Inference from Iterative Simulation Using Multiple Sequences. Statist. Sci. 7:457–472.

Grabherr MG, Haas BJ, Yassour M, Levin JZ, Thompson DA, Amit I, Adiconis X, Fan L, Raychowdhury R, Zeng Q, et al. 2011. Full-length transcriptome assembly from RNA-Seq data without a reference genome. Nature Biotechnology 29:644–652.

Gurney, R. 1942. Larvae of Decapod Crustacea. Ray Society, 129:1–306.

Haas BJ, Papanicolaou A, Yassour M, Grabherr M, Blood PD, Bowden J, Couger MB, Eccles D, Li B, Lieber M, et al. 2013. De novo transcript sequence reconstruction from RNA-seq using the Trinity platform for reference generation and analysis. Nature Protocols 8:1494–1512.

Heath TA, Hedtke SM, Hillis DM. 2008. Taxon sampling and the accuracy of phylogenetic analyses. Journal of Systematics and Evolution 46:239–257.

Hedtke SM, Townsend TM, Hillis DM. 2006. Resolution of Phylogenetic Conflict in Large Data Sets by Increased Taxon Sampling. Systematic Biology 55:522–529.

Hegna TA, Luque J, Wolfe JM. 2020. The fossil record of the Pancrustacea. In: Poore Gcb, Thiel M, editors. Evolution and Biogeography. Vol. 8. Natural History of the Crustacea. Oxford: Oxford University Press. p. 21–52.

Hellmuth M, Wieseke N, Lechner M, Lenhof H-P, Middendorf M, Stadler PF. 2015. Phylogenomics with paralogs. Proceedings of the National Academy of Sciences 112:2058–2063.

Hendy MD, Penny D. 1989. A Framework for the Quantitative Study of Evolutionary Trees. Systematic Zoology 38:297.

Hernandez AM, Ryan JF. 2021. Six-State Amino Acid Recoding is not an Effective Strategy to Offset Compositional Heterogeneity and Saturation in Phylogenetic Analyses. Systematic Biology 70:1200–1212.

Höpel CG, Yeo D, Grams M, Meier R, Richter S. 2022. Mitogenomics supports the monophyly of Mysidacea and Peracarida (Malacostraca). Zoologica Scripta:zsc.12554.

Hu F, Lin Y, Tang J. 2014. MLGO: phylogeny reconstruction and ancestral inference from gene-order data. BMC Bioinformatics 15:354.

Huang H, He Q, Kubatko LS, Knowles LL. 2010. Sources of Error Inherent in Species-Tree Estimation: Impact of Mutational and Coalescent Effects on Accuracy and Implications for Choosing among Different Methods. Systematic Biology 59:573–583.

Huelsenbeck JP, Larget B, Miller RE, Ronquist F. 2002. Potential Applications and Pitfalls of Bayesian Inference of Phylogeny. Systematic Biology 51:673–688.

Huerta-Cepas J, Serra F, Bork P. 2016. ETE 3: Reconstruction, Analysis, and Visualization of Phylogenomic Data. Mol Biol Evol 33:1635–1638.

Hugall AF, Lee MSY. 2007. The Likelihood Node Density Effect and Consequences for Evolutionary Studies of Molecular Rates. Evolution 61:2293–2307.

Huys R, Boxshall GA. 1991. Copepod Evolution. London: Ray Society Available from: http://193.190.8.15/dpm/handle/0/3577

Huys R, Boxshall GA, Lincoln RJ. 1993. The Tantulocaridan Life Cycle: the Circle Closed? Journal of Crustacean Biology 13:432–442.

Jombart T, Kendall M, Almagro-Garcia J, Colijn C. 2017. treespace: Statistical exploration of landscapes of phylogenetic trees. Mol Ecol Resour 17:1385–1392.

Katoh K, Standley DM. 2013. MAFFT Multiple Sequence Alignment Software Version 7: Improvements in Performance and Usability. Molecular Biology and Evolution 30:772–780.

Kendall M, Colijn C. 2016. Mapping Phylogenetic Trees to Reveal Distinct Patterns of Evolution. Mol Biol Evol 33:2735–2743.

Kishino, H, Miyata, T, Hasegawa, M. 1990. Maximum likelihood inference of protein phylogeny and the origin of chloroplasts. J Mol Evol 31:151–160. https://doi.org/10.1007/BF02109483

Koga C, Rouse GW. 2021. Mitogenomics and the Phylogeny of Mantis Shrimps (Crustacea: Stomatopoda). Diversity 13:647.

Lanfear R, Calcott B, Kainer D, Mayer C, Stamatakis A. 2014. Selecting optimal partitioning schemes for phylogenomic datasets. BMC Evol Biol 14:82.

Lanfear R, Frandsen PB, Wright AM, Senfeld T, Calcott B. 2016. PartitionFinder 2: New Methods for Selecting Partitioned Models of Evolution for Molecular and Morphological Phylogenetic Analyses. Molecular Biology and Evolution 34:msw260.

Lanier HC, Knowles LL. 2015. Applying species-tree analyses to deep phylogenetic histories: Challenges and potential suggested from a survey of empirical phylogenetic studies. Molecular Phylogenetics and Evolution 83:191–199.

Lartillot N, Brinkmann H, Philippe H. 2007. Suppression of long-branch attraction artefacts in the animal phylogeny using a site-heterogeneous model. BMC Evol Biol 7:S4.

Lartillot N, Philippe H. 2004. A Bayesian Mixture Model for Across-Site Heterogeneities in the Amino-Acid Replacement Process. Molecular Biology and Evolution 21:1095–1109.

Lartillot N, Rodrigue N, Stubbs D, Richer J. 2013. PhyloBayes MPI: Phylogenetic Reconstruction with Infinite Mixtures of Profiles in a Parallel Environment. Systematic Biology 62:611–615.

Laumer CE, Fernández R, Lemer S, Combosch D, Kocot KM, Riesgo A, Andrade SCS, Sterrer W, Sørensen MV, Giribet G. 2019. Revisiting metazoan phylogeny with genomic sampling of all phyla. Proc. R. Soc. B 286:20190831.

Le SQ, Gascuel O, Lartillot N. 2008. Empirical profile mixture models for phylogenetic reconstruction. Bioinformatics 24:2317–2323.

Lee MSY, Soubrier J, Edgecombe GD. 2013. Rates of Phenotypic and Genomic Evolution during the Cambrian Explosion. Current Biology 23:1889–1895.

Lepage T, Bryant D, Philippe H, Lartillot N. 2007. A General Comparison of Relaxed Molecular Clock Models. Molecular Biology and Evolution 24:2669–2680.

Lewin HA, Richards S, Lieberman Aiden E, Allende ML, Archibald JM, Bálint M, Barker KB, Baumgartner B, Belov K, Bertorelle G, et al. 2022. The Earth BioGenome Project 2020: Starting the clock. Proc. Natl. Acad. Sci. U.S.A. 119:e2115635118.

Lewin HA, Robinson GE, Kress WJ, Baker WJ, Coddington J, Crandall KA, Durbin R, Edwards SV, Forest F, Gilbert MTP, et al. 2018. Earth BioGenome Project: Sequencing life for the future of life. Proc Natl Acad Sci USA 115:4325–4333.

Li W, Godzik A. 2006. Cd-hit: a fast program for clustering and comparing large sets of protein or nucleotide sequences. Bioinformatics 22:1658–1659.

Li Y, Shen X-X, Evans B, Dunn CW, Rokas A. 2021. Rooting the animal tree of life. Molecular Biology and Evolution 38:4322–4333.

Lozano-Fernandez J, Carton R, Tanner AR, Puttick MN, Blaxter M, Vinther J, Olesen J, Giribet G, Edgecombe GD, Pisani D. 2016. A molecular palaeobiological exploration of arthropod terrestrialization. Philosophical Transactions of the Royal Society B: Biological Sciences 371:20150133.

Lozano-Fernandez J, Giacomelli M, Fleming J, Chen A, Vinther J, Thomsen PF, Glenner H, Palero F, Legg DA, Iliffe TM, et al. 2019. Pancrustacean evolution illuminated by taxon-rich genomic-scale data sets with an expanded remipede sampling. Genome Biology and Evolution 11:2055–2070.

Lozano-Fernandez J, Tanner AR, Puttick MN, Vinther J, Edgecombe GD, Pisani D. 2020. A Cambrian– Ordovician Terrestrialization of Arachnids. Front. Genet. 11:182.

Luque J, Gerken S. 2019. Exceptional preservation of comma shrimp from a mid-Cretaceous Lagerstätte of Colombia, and the origins of crown Cumacea. Proc. R. Soc. B 286:20191863.

Mallatt J, Giribet G. 2006. Further use of nearly complete 28S and 18S rRNA genes to classify Ecdysozoa: 37 more arthropods and a kinorhynch. Molecular Phylogenetics and Evolution 40:772–794.

McClain CR, Balk MA, Benfield MC, Branch TA, Chen C, Cosgrove J, Dove ADM, Gaskins L, Helm RR, Hochberg FG, et al. 2015. Sizing ocean giants: patterns of intraspecific size variation in marine megafauna. PeerJ 3:e715.

Mirarab S, Bayzid MS, Boussau B, Warnow T. 2014. Statistical binning enables an accurate coalescent-based estimation of the avian tree. Science 346:1250463–1250463.

Mirarab S, Bayzid MS, Warnow T. 2016. Evaluating Summary Methods for Multilocus Species Tree Estimation in the Presence of Incomplete Lineage Sorting. Syst Biol 65:366–380.

Mirarab S, Warnow T. 2015. ASTRAL-II: coalescent-based species tree estimation with many hundreds of taxa and thousands of genes. Bioinformatics 31:i44–i52.

Møller OS, Olesen J, Avenant-Oldewage A, Thomsen PF, Glenner H. 2008. First maxillae suction discs in Branchiura (Crustacea): Development and evolution in light of the first molecular phylogeny of Branchiura, Pentastomida, and other “Maxillopoda.” Arthropod Structure & Development 37:333–346.

Mongiardino Koch NM. 2021. Phylogenomic subsampling and the search for phylogenetically reliable loci. Molecular Biology and Evolution 38:4025–4038.

Moret BME, Warnow T. 2005. Advances in phylogeny reconstruction from gene order and content data. Methods in Enzymology 395:673–700.

Nabhan AR, Sarkar IN. 2012. The impact of taxon sampling on phylogenetic inference: a review of two decades of controversy. Briefings in Bioinformatics 13:122–134.

Nguyen L-T, Schmidt HA, von Haeseler A, Minh BQ. 2015. IQ-TREE: A Fast and Effective Stochastic Algorithm for Estimating Maximum-Likelihood Phylogenies. Molecular Biology and Evolution 32:268–274.

Oakley TH, Wolfe JM, Lindgren AR, Zaharoff AK. 2013. Phylotranscriptomics to Bring the Understudied into the Fold: Monophyletic Ostracoda, Fossil Placement, and Pancrustacean Phylogeny. Molecular Biology and Evolution 30:215–233.

One Thousand Plant Transcriptomes Initiative. 2019. One thousand plant transcriptomes and the phylogenomics of green plants. Nature 574:679–685.

Owen CL, Stern DB, Hilton SK, Crandall KA. 2020. Hemiptera phylogenomic resources: Tree□based orthology prediction and conserved exon identification. Mol Ecol Resour 20:1346–1360.

Patel S, Kimball RT, Braun EL. 2013. Error in Phylogenetic Estimation for Bushes in the Tree of Life. Journal of Phylogenetics & Evolutionary Biology 01:110.

Petrunina AS, Høeg JT, Kolbasov GA. 2018. Anatomy of the Tantulocarida: first results obtained using TEM and CLSM. Part I: tantulus larva. Org Divers Evol 18:459–477.

Petrunina AS, Neretina TV, Mugue NS, Kolbasov GA. 2014. Tantulocarida versus Thecostraca: inside or outside? First attempts to resolve phylogenetic position of Tantulocarida using gene sequences. Journal of Zoological Systematics and Evolutionary Research 52:100–108.

Poe S. 2003. Evaluation of the Strategy of Long-Branch Subdivision to Improve the Accuracy of Phylogenetic Methods. Systematic Biology 52:423–428.

Price MN, Dehal PS, Arkin AP. 2010. FastTree 2–approximately maximum-likelihood trees for large alignments. PloS one 5:e9490.

Puslednik L, Serb JM. 2008. Molecular phylogenetics of the Pectinidae (Mollusca: Bivalvia) and effect of increased taxon sampling and outgroup selection on tree topology. Molecular Phylogenetics and Evolution 48:1178–1188.

Puttick MN. 2019. MCMCtreeR: functions to prepare MCMCtree analyses and visualise posterior ages on trees. Bioinformatics:3.

Regier J, Shultz J, Ganley A, Hussey A, Shi D, Ball B, Zwick A, Stajich J, Cummings M, Martin J, et al. 2008. Resolving Arthropod Phylogeny: Exploring Phylogenetic Signal within 41 kb of Protein-Coding Nuclear Gene Sequence. Systematic Biology 57:920–938.

Regier JC, Shultz JW, Kambic RE. 2005. Pancrustacean phylogeny: hexapods are terrestrial crustaceans and maxillopods are not monophyletic. Proceedings of the Royal Society B: Biological Sciences 272:395–401.

Regier JC, Shultz JW, Zwick A, Hussey A, Ball B, Wetzer R, Martin JW, Cunningham CW. 2010. Arthropod relationships revealed by phylogenomic analysis of nuclear protein-coding sequences. Nature 463:1079–1083.

dos Reis M, Donoghue PCJ, Yang Z. 2015. Bayesian molecular clock dating of species divergences in the genomics era. Nature Reviews Genetics 17:71–80.

dos Reis M, Thawornwattana Y, Angelis K, Telford MJ, Donoghue PCJ, Yang Z. 2015. Uncertainty in the Timing of Origin of Animals and the Limits of Precision in Molecular Timescales. Current Biology 25:2939–2950.

dos Reis M, Yang Z. 2011. Approximate Likelihood Calculation on a Phylogeny for Bayesian Estimation of Divergence Times. Molecular Biology and Evolution 28:2161–2172.

von Reumont BM, Jenner RA, Wills MA, Dell’Ampio E, Pass G, Ebersberger I, Meyer B, Koenemann S, Iliffe TM, Stamatakis A, et al. 2012. Pancrustacean Phylogeny in the Light of New Phylogenomic Data: Support for Remipedia as the Possible Sister Group of Hexapoda. Molecular Biology and Evolution 29:1031–1045.

Richter S, Scholtz G. 2001. Phylogenetic analysis of the Malacostraca (Crustacea). Journal of Zoological Systematics and Evolutionary Research 39:113–136.

Robin N, Gueriau P, Luque J, Jarvis D, Daley AC, Vonk R. 2021. The oldest peracarid crustacean reveals a Late Devonian freshwater colonization by isopod relatives. Biol. Lett. 17:20210226.

Rokas A, Carroll SB. 2006. Bushes in the Tree of Life. PLoS Biol 4:e352.

Roskov Y, Ower G, Orrell T, Nicolson D, Bailly N, Kirk PM, Bourgoin T, DeWalt RE, Decock W, Nieukerken E van, Zarucchi J, Penev L, editors. 2022. Species 2000 & ITIS Catalogue of Life, 10th February 2022. Digital resource at https://www.catalogueoflife.org/col. Species 2000: Naturalis, Leiden, the Netherlands. ISSN 2405-8858.

Rota-Stabelli O, Campbell L, Brinkmann H, Edgecombe GD, Longhorn SJ, Peterson KJ, Pisani D, Philippe H, Telford MJ. 2011. A congruent solution to arthropod phylogeny: phylogenomics, microRNAs and morphology support monophyletic Mandibulata. Proceedings of the Royal Society B: Biological Sciences 278:298–306.

Rota-Stabelli O, Daley AC, Pisani D. 2013. Molecular timetrees reveal a Cambrian colonization of land and a new scenario for ecdysozoan evolution. Current Biology 23:392–398.

Rota-Stabelli O, Lartillot N, Philippe H, Pisani D. 2013. Serine codon-usage bias in deep phylogenomics: Pancrustacean relationships as a case study. Systematic Biology 62:121–133.

Sayyari E, Mirarab S. 2016. Fast Coalescent-Based Computation of Local Branch Support from Quartet Frequencies. Mol Biol Evol 33:1654–1668.

Schram FR. 1984. Fossil Syncarida. Transactions of the San Diego Society of Natural History 20:189–246.

Schram FR, Hof CH. 1998. Fossils and the interrelationships of major crustacean groups. In: Edgecombe GD, editor. Arthropod fossils and phylogeny. New York: Columbia University Press. p. 233–302.

Schwentner M, Combosch DJ, Pakes Nelson J, Giribet G. 2017. A Phylogenomic Solution to the Origin of Insects by Resolving Crustacean-Hexapod Relationships. Current Biology 27:1818-1824.e5.

Schwentner M, Richter S, Rogers DC, Giribet G. 2018. Tetraconatan phylogeny with special focus on Malacostraca and Branchiopoda: highlighting the strength of taxon-specific matrices in phylogenomics. Proceedings of the Royal Society B: Biological Sciences 285:20181524.

Serban, E, 1972. Bathynella (Podophallocarida, Bathynellacea). Travaux de l’Institut de Spéologie ‘Émile Racovitza’ 11:11–225.

Serban, E, 1973. Sur le processus de la pléonisation du péréion dans l’ordre des bathynellacea (Crustacea, Malacostraca, Podophallocarida). Bijdragen tot de Dierkunde 43:173–201.

Shen X-X, Hittinger CT, Rokas A. 2017. Contentious relationships in phylogenomic studies can be driven by a handful of genes. Nature Ecology & Evolution 1:0126.

Shen X-X, Salichos L, Rokas A. 2016. A genome-scale investigation of how sequence-, function-, and tree-based gene properties influence phylogenetic inference. Genome Biology and Evolution 8:2565–2580.

Shimodaira H. 2002. An Approximately Unbiased Test of Phylogenetic Tree Selection. Systematic Biology 51:492–508.

Siewing R. 1956. Untersuchungen zur Morphologie der Malacostraca (Crustacea). Zoologische Jahrbücher, Abteilung Anatomie und Ontogenie der Tiere 75:39–176.

Simakov O, Bredeson J, Berkoff K, Marletaz F, Mitros T, Schultz DT, O’Connell BL, Dear P, Martinez DE, Steele RE, et al. 2022. Deeply conserved synteny and the evolution of metazoan chromosomes. Sci. Adv. 8:eabi5884.

Smith ML, Hahn MW. 2021. New Approaches for Inferring Phylogenies in the Presence of Paralogs. Trends in Genetics 37:174–187.

Smith ML, Hahn MW. 2022. The Frequency and Topology of Pseudoorthologs. Systematic Biology 71:649–659.

Smith ML, Vanderpool D, Hahn MW. 2022. Using all Gene Families Vastly Expands Data Available for Phylogenomic Inference. Molecular Biology and Evolution 39:msac112.

Smith SA, Dunn CW. 2008. Phyutility: a phyloinformatics tool for trees, alignments and molecular data. Bioinformatics 24:715–716.

Smith SA, Pease JB. 2017. Heterogeneous molecular processes among the causes of how sequence similarity scores can fail to recapitulate phylogeny. Briefings in Bioinformatics 18:451–457.

Smith SA, Wilson NG, Goetz FE, Feehery C, Andrade SCS, Rouse GW, Giribet G, Dunn CW. 2011. Resolving the evolutionary relationships of molluscs with phylogenomic tools. Nature 480:364–367.

Song L, Florea L. 2015. Rcorrector: efficient and accurate error correction for Illumina RNA-seq reads. GigaSci 4:48.

Stamatakis A. 2014. RAxML version 8: a tool for phylogenetic analysis and post-analysis of large phylogenies. Bioinformatics 30:1312–1313.

Steenwyk JL, Buida TJ, Labella AL, Li Y, Shen X-X, Rokas A. 2021. PhyKIT: a broadly applicable UNIX shell toolkit for processing and analyzing phylogenomic data. Bioinformatics 37:2325–2331.

Strimmer, K, Rambaut, A. 2002. Inferring confidence sets of possibly misspecified gene trees. Proceedings of the Royal Society of London. Series B: Biological Sciences 269: 137–142.

Struck T. 2014. TreSpEx--Detection of Misleading Signal in Phylogenetic Reconstructions Based on Tree Information. Evolutionary Bioinformatics 10:EBO–S14239.

Susko E, Roger AJ. 2007. On Reduced Amino Acid Alphabets for Phylogenetic Inference. Molecular Biology and Evolution 24:2139–2150.

Talavera G, Castresana J. 2007. Improvement of Phylogenies after Removing Divergent and Ambiguously Aligned Blocks from Protein Sequence Alignments. Systematic Biology 56:564–577.

Van Dongen S. 2000. Graph Clustering by Flow Simulation. PhD thesis, University of Utrecht.

Van Dongen S. 2008. Graph clustering via a discrete uncoupling process. SIAM Journal on Matrix Analysis and Applications 30:121–141.

Wang H-C, Minh BQ, Susko E, Roger AJ. 2018. Modeling Site Heterogeneity with Posterior Mean Site Frequency Profiles Accelerates Accurate Phylogenomic Estimation. Systematic Biology 67:216–235.

Warnock RCM, Yang Z, Donoghue PCJ. 2012. Exploring uncertainty in the calibration of the molecular clock. Biology Letters 8:156–159.

Weigert A, Helm C, Meyer M, Nickel B, Arendt D, Hausdorf B, Santos SR, Halanych KM, Purschke G, Bleidorn C, et al. 2014. Illuminating the Base of the Annelid Tree Using Transcriptomics. Molecular Biology and Evolution 31:1391–1401.

Whelan NV, Halanych KM. 2017. Who Let the CAT Out of the Bag? Accurately Dealing with Substitutional Heterogeneity in Phylogenomic Analyses. Systematic Biology 66:232–255.

Whitfield JB, Lockhart PJ. 2007. Deciphering ancient rapid radiations. Trends in Ecology & Evolution 22:258–265.

Wolfe JM. 2017. Metamorphosis is ancestral for crown euarthropods, and evolved in the Cambrian or earlier. Integrative and Comparative Biology 57:499–509.

Wolfe JM, Breinholt JW, Crandall KA, Lemmon AR, Moriarty Lemmon E, Timm LE, Siddall ME, Bracken-Grissom HD. 2019. A phylogenomic framework, evolutionary timeline, and genomic resources for comparative studies of decapod crustaceans. Proceedings of the Royal Society B: Biological Sciences 286:20190079.

Wolfe JM, Daley AC, Legg DA, Edgecombe GD. 2016. Fossil calibrations for the arthropod Tree of Life. Earth-Science Reviews 160:43–110.

WoRMS (2022). Crustacea. Accessed at: https://www.marinespecies.org/aphia.php?p=taxdetails&id=1066 on 2022-09-22

Xi Z, Liu L, Davis CC. 2015. Genes with minimal phylogenetic information are problematic for coalescent analyses when gene tree estimation is biased. Molecular Phylogenetics and Evolution 92:63–71.

Yang Y, Smith SA. 2014. Orthology Inference in Nonmodel Organisms Using Transcriptomes and Low-Coverage Genomes: Improving Accuracy and Matrix Occupancy for Phylogenomics. Molecular Biology and Evolution 31:3081–3092.

Zhai D, Ortega-Hernández J, Wolfe JM, Hou X, Cao C, Liu Y. 2019. Three-dimensionally preserved appendages in an early Cambrian stem-group pancrustacean. Current Biology 29:171–177.

Zhang C, Rabiee M, Sayyari E, Mirarab S. 2018. ASTRAL-III: polynomial time species tree reconstruction from partially resolved gene trees. BMC Bioinformatics 19:153.

Zhang X, Siveter DJ, Waloszek D, Maas A. 2007. An epipodite-bearing crown-group crustacean from the Lower Cambrian. Nature 449:595–598.

