## Supplementary material for "Major revisions in pancrustacean phylogeny with recommendations for resolving challenging nodes": Table S7

**Supplement 7. Fossil dates and justifications**

**Table S7.** Fossil calibrations.

| **Node** | **Fossil taxon** | **Minimum age (Ma)** | **Soft maximum age (Ma)** |
| --- | --- | --- | --- |
| Root | *Yicaris dianensis* | 514 | 636.1 |
| Myodocopa | *Luprisca incuba* | 443.8 | 509 |
| Branchiopoda | *Lepidocaris rhyniensis* | 405 | 521 |
| Collembola | *Rhyniella praecursor* | 405 | 521 |
| Microcoryphia | *Gigamachilis triassicus* | 239.36 | 521 |
| Holometabola | *Westphalomerope maryvonneae* | 313.70 | 411 |
| Cirripedia | *Illilepas damrowi* | 306.9 | 521 |
| Verunipeltata | *Ursquilla yehoachi* | 79.01 | 521 |
| Dendrobranchiata | *Ifasya madagascariensis* | 251.2 | 521 |
| Reptantia | *Palaeopalaemon newberryi* | 358.5 | 521 |
| Peracarida | *Oxyuropoda ligioides* | 358.2 | 521 |
| Talitroida | *Caecorchestia bousfieldi* | 23.0 | 365.7 |
| Cumacea | *Eobodotria muisca* | 93.9 | 521 |

**Fossil Calibration Justifications:**

1. ***Node*.** This node comprises crown group Myodocopa. In our phylogeny, this is the clade comprising Halocyprida and Myodocopida, their last common ancestor and all of its descendants. Monophyly has been previously established by phylogenetic analysis of combined morphological and transcriptomic data (Oakley et al., 2013) and transcriptomes alone (Schwentner et al., 2018). All calibration data as in Wolfe et al. (2016), node 39.
2. ***Node*.** This node comprises crown group Branchiopoda. In our phylogeny, this is the clade comprising Anostraca, Notostraca, and Cladocera, their last common ancestor and all of its descendants. Monophyly has been previously established by phylogenetic analysis of combined morphological and transcriptomic data (Oakley et al. 2013) and transcriptomes alone (Lozano-Fernandez et al., 2019; Schwentner et al., 2018). All calibration data as in Wolfe et al. (2016), node 58.
3. ***Node*.** This node comprises crown group Collembola. In our phylogeny, this is the clade comprising Entomobryomorpha, Poduromorpha and Symphypleona, their last common ancestor and all of its descendants. All calibration data as in Wolfe et al. (2016), node 65.
4. ***Node*.** This node comprises crown group Microcoryphia (the crown group of Archaeognatha). This clade comprises the families ‘Machilidae’ and Meinertillidae, their last common ancestor and all of its descendants. Further discussion in Wolfe et al. (2016), node 68.

***Fossil specimens.*** *Gigamachilis triassicus* Montagna et al., 2017. MCSN8463 (Museo Cantonale di Storia Naturale di Lugano, Switzerland), holotype.

***Phylogenetic justification.*** The *G. triassicus* holotype is exceptional, preserving the central nervous system and ventral nerve cord as phosphate replicates (Montagna et al., 2017), although these morphologies are not informative about assignment within Microcoryphia. The external morphologies of the fossil, such as large maxillary palps, abdominal coxopodal vesicles and styli, and structures of the cerci, all assign *G. triassicus* to ‘Machilidae’ (Montagna et al., 2017, 2019). While ‘Machilidae’ are likely paraphyletic, revised morphological phylogenetic analyses with both maximum parsimony and Bayesian inference recover *G. triassicus* within Machilidae *sensu stricto* (Montagna 2020), and therefore within the crown group of Microcoryphia.

***Age justification.*** The fossil of *G. triassicus* was found at locality/site D on the uppermost Lower Kalkschieferzone of the Monte San Giorgio Lagerstätte, Switzerland (Montagna et al. 2017). Site D has been radiometrically dated using U-Pb dating, retrieving an age of 239.51 Ma ± 0.15 Myr (Stockar et al. 2012). The previous work using *G. triassicus* as a calibration (Montagna et al. 2019) erroneously did not include the measurement error, so we use a slightly different minimum age of 239.36 Ma.

Soft maximum as in Wolfe et al. (2016), node 68.

5. ***Node*.** This node comprises crown group Holometabola. In our phylogeny, this is the clade comprising Hymenoptera, Coleoptera, Lepidoptera, and Diptera, their last common ancestor and all of its descendants. All calibration data as in Wolfe et al. (2016), node 89.
6. ***Node*.** This node comprises crown group Cirripedia. In our phylogeny, this is the clade comprising Rhizocephala and Thoracica, their last common ancestor and all of its descendants. Monophyly has been established in several individual phylogenetic analyses of molecular and morphological data, and recently using a synthesis approach (Ewers-Saucedo et al., 2019) and taxonomic revision (Chan et al., 2021). Acrothoracica are also included in Cirripedia, but were not sampled in our phylogeny. All calibration data remain the same as in Wolfe et al. (2016), node 45.
7. ***Node.*** This node comprises crown group Verunipeltata (the crown group of Stomatopoda). In our phylogeny, this is the clade comprising Lysiosquilloidea, Squilloidea, and Gonodactyloidea, their last common ancestor and all of its descendants. All calibration data as in Wolfe et al. (2016), node 51. Although our phylogeny recovers paraphyly of the genus *Neogonodactylus*, the whole clade and fossil position are unaffected.
8. ***Node*.** This node comprises crown group Dendrobranchiata. In our phylogeny, this is the clade comprising Sergestoidea and Penaeoidea, their last common ancestor and all of its descendants. Monophyly has been previously established by phylogenetic analysis of target capture data (Wolfe et al. 2019). All calibration data as in Wolfe et al. (2019), node 3. Note that the node calibrated previously was Penaeoidea, but total evidence analyses (including fossils) do not uncover a stem group penaeoid (Robalino et al., 2016), therefore the calibration for the superfamily also applies to Dendrobranchiata.
9. ***Node*.** This node comprises crown group Reptantia. In our phylogeny, this is the clade comprising Astacidea, Anomura, and Brachyura, their last common ancestor and all of its descendants. Monophyly has been previously established by phylogenetic analysis of target capture data (Wolfe et al. 2019) and transcriptomics (Schwentner et al. 2018). All calibration data as in Wolfe et al. (2019), node 6.
10. ***Node*.** This node comprises crown group Peracarida. In our phylogeny, this is the clade comprising Stygiomysida, Mysida, Tanaidacea, Cumacea, Isopoda, and Amphipoda, their last common ancestor and all of its descendants. Although some studies have questioned the relationship of Mysida to Peracarida, our analysis and a previous transcriptomic analysis (Schwentner et al. 2018) support their inclusion.

***Fossil specimens*.** *Oxyuropoda ligioides* Carpenter & Swain 1908. NMING (National Museum of Ireland, Dublin, Natural History) F7633, holotype. Redescribed by Robin et al. (2021).

***Phylogenetic justification*.** In the redescription of *O. ligioides*, a Bayesian morphological phylogeny recovered the fossil as sister group to the clade containing Amphipoda, Isopoda, Cumacea, and Tanaidacea (Robin et al., 2021). While this is not the topology that we recovered for the extant clades of peracarids, the morphology of *O. ligioides* supports its position within crown group Peracarida.

***Age justification*.** The fossil was collected at the Classic or Old Quarry of Kiltorcan Hill, County Kilkenny, Ireland (Robin et al., 2021). The Old Quarry is no longer available for stratigraphic study (Jarvis 2000), but a minimum age can be assessed based on the overlying Roadstone Quarry. The fossiliferous layer of the Roadstone Quarry yields a combination of palynological material indicative of the VI Miospore Biozone (Jarvis 1990). The VI Miospore Biozone in Ireland is correlated to several *Siphonodella* conodont Biozones globally, with the uppermost being the *Siphonodella jii* Biozone (Graham and Sevastopulo 2021) at approximately 358.2 Ma (Aretz et al. 2020). While Robin et al. (2021) estimated a latest Devonian age for the Old Quarry, we conservatively adopt the minimum age of 358.2 Ma. Soft maximum as in Wolfe et al. (2016), node 52.

***Discussion.*** In Wolfe et al. (2016), this node was calibrated by *Hesslerella shermani*, a crown group phreatoicidean isopod. Additional discussion of the early peracarid record in Hegna et al. (2020b) had already suggested older calibrations may be possible but required a phylogenetic context. However, the phylogenetic analysis of Robin et al. (2021) also incorporated one of the candidate taxa, *Tealliocaris*, which was not recovered within peracarids.

11. ***Node*.** This node comprises crown group Talitroida. In our phylogeny, this is the clade comprising Hyalellidae and Talitridae, their last common ancestor and all of its descendants. Monophyly of this clade has been found in a recent multilocus molecular phylogeny (Copilaş-Ciocianu et al., 2019a). Apart from recent efforts focused on transcriptomes of Lake Baikal gammaroids (Naumenko et al., 2016), multilocus data in crangonyctids and haustoriids (Copilaş-Ciocianu et al., 2019b; Hancock et al., 2020), and morphological analysis at an exemplar level (Lowry and Myers, 2017), broader scale amphipod phylogeny remains relatively poorly studied.

***Fossil specimens*.** *Caecorchestia bousfieldi* Hegna & Lazo-Wasem 2020. IHNFG-5824 (Museo de Paleontología 'Eliseo Palacios Aguilera', Tuxtla Gutiérrez, Chiapas, Mexico), holotype (Hegna et al., 2020a).

***Phylogenetic justification*.** The description of *C. bousfieldi* followed the diagnosis of Lowry & Myers (2013) for membership in Talitridae. No suggestions were made as to how *C. bousfieldi* relates to particular genera of Talitridae (Hegna et al. 2020a), however, even a position on the stem lineage of the family would remain an appropriate calibration for Talitroida.

***Age justification*.** This fossil was discovered in amber from Campo La Granja mine, near Simojovel, Chiapas, Mexico (Hegna et al. 2020a). The amber bearing layer at this locality is part of the Finca Carmitto Member of the La Quinta Formation, which underlies the Mazantic Shale (Serrano-Sánchez et al. 2015). In the absence of an additional radiometric age for the La Quinta Formation, we adopt the minimum age as for the lower Mazantic Shale, while acknowledging that the amber itself may have different ages, as follows. Benthic foraminiferal fossils indicate the Mazantic Shale belongs to the *Siphogenerina transversa* regional Zones (correlated to planktonic foram Zones N8-N9 and M5-M6; Solórzano Kraemer, 2007). The global Zones N8-N9 and M5-M6 represent a Langhian age (Serrano-Sánchez et al. 2015) with a correlation to the C5AD chron that has a minimum age of 14.609 Ma (Hilgen et al., 2012). However, 87Sr/86Sr radiometric dates from the lower Mazantic Shale may be precisely at the Oligocene-Miocene boundary (Vega et al. 2009) with an age of 23.0 Ma. One possibility to reconcile these dates is that the amber fossils themselves may be somewhat reworked relative to the sediment. Although it is a less conservative choice, we currently suggest the Mazantic Shale radiometric age provides a minimum of 23.0 Ma.

A soft maximum age is assessed based on the assumption that the oldest Talitroida would be younger than the oldest Peracarida, which is *O. ligioides* (node 11 herein). The maximum age of the Old Quarry deposit of Kiltorcan Hill, Ireland, can only be constrained by the older New Quarry (Robin et al. 2012, Jarvis 1990). Palynological material at the New Quarry correlates to the LE Miospore Biozone, and therefore the *Siphonodella praesulcata* conodont Biozone (Jarvis 1990; Graham and Sevastopulo 2021). Radiometric dating provides an estimate of 363.3 Ma ± 2.4 Myr for the *S. praesulcata* maximum (Becker et al. 2012), and therefore a soft maximum age of 365.7 Ma.

***Discussion.*** A number of other amphipod fossils are known from older Baltic amber (reviewed by Hegna et al. 2020a, some presented as calibrations by Copilaş-Ciocianu et al. 2019a; Hancock et al. 2020). However, none of these taxa can be clearly assigned to a clade found in our transcriptome phylogeny. Calibrations have also been presented based on biogeographic distribution (e.g. Hancock et al. 2020), but these are too shallow to be captured by our study.

12. ***Node*.** This node comprises crown group Cumacea. In our phylogeny, this is the clade comprising Diastylidae and Nannastacidae, their last common ancestor and all of its descendants. Monophyly is established in a phylogenetic analysis of morphological data (Luque and Gerken, 2019), and represents deep splits within crown group Cumacea. No molecular analysis has adequately tested cumacean monophyly.

***Fossil specimens*.** *Eobodotria muisca* Luque & Gerken 2019. IGM (Colombian Geological Survey, Bogotá, Colombia) p880694a (holotype, male) and IGM p880691a (paratype, female).

***Phylogenetic justification*.** Based on a morphological phylogenetic analysis, *E. musica* is within the crown group family Bodotriidae (Luque & Gerken 2019). As Bodotriidae resolves in a polytomy with Nannastacidae to the exclusion of several other families (including Diastylidae), the fossil is certainly within the crown group.

***Age justification*.** These fossils were collected in a recently described Lagerstätte belonging to the Churuvita Group, near Pesca, Department of Boyacá, Eastern Cordillera of Colombia (Luque & Gerken 2019; Luque et al. 2019). *E. muisca* is known from the lowermost segment A of the Churuvita Formation, which bears few index fossils, while the overlying segment B is not fossiliferous (Luque et al., 2019). Therefore, we constrain the age from the base of uppermost segment C, which is part of the San Rafael Formation. *Watinoceras* and *Hoplitoides* ammonite (Feldmann et al., 1999), *Mytiloides* bivalve, and foraminiferal (Sánchez-Quiñónez and Tchegliakova, 2005) index fossils constrain the base of the San Rafael Formation to the lower Turonian. More precise global zonation is not available. Therefore, the minimum age of the Churuvita Formation, at its boundary with the overlying San Rafael Formation, is estimated at 93.9 Ma (the global boundary between Cenomanian and Turonian stages). The soft maximum age follows Wolfe et al. (2016), node 52.

**References:**

Aretz, M., Herbig, H.G., Wang, X.D., Gradstein, F.M., Agterberg, F.P. , Ogg, J.G., 2020. The Carboniferous Period, in: Gradstein, F.M.,Ogg, J.G., Schmitz, M.D., Ogg., G.M. (Eds.), Geologic Time Scale 2020. Elsevier, pp. 811-874.

Becker, R.T., Gradstein, F.M., Hammer, O., 2012. The Devonian Period, in: The Geologic Time Scale. Elsevier, pp. 559-601.

Carpenter, G.H., Swain, I., 1908. A new Devonian isopod from Kiltorcan, County Kilkenny. Proc. R. Irish Acad. B 27, 61–67.

Chan, B.K.K., Dreyer, N., Gale., A.S., Glenner, H., Ewers-Saucedo, C., Pérez-Losada, M., Crandall, K.A., Høeg, J.T., 2021. The evolutionary diversity of barnacles, with an updated classification of fossil and living forms. Zool. J. Linn. Soc. <https://doi.org/10.1093/zoolinnean/zlaa160>

Copilaş-Ciocianu, D., Borko, Š., Fišer, C., 2019a. The late blooming amphipods: global change promoted post-Jurassic ecological radiation despite Palaeozoic origin. Mol. Phylogenet. Evol. <https://doi.org/10.1101/675140>

Copilaş-Ciocianu, D., Sidorov, D., Gontcharov, A., 2019b. Adrift across tectonic plates: molecular phylogenetics supports the ancient Laurasian origin of old limnic crangonyctid amphipods. Org. Divers. Evol. <https://doi.org/10.1007/s13127-019-00401-7>

Ewers-Saucedo, C., Owen, C.L., Pérez-Losada, M., Høeg, J.T., Glenner, H., Chan, B.K.K., Crandall, K.A., 2019. Towards a barnacle tree of life: integrating diverse phylogenetic efforts into a comprehensive hypothesis of thecostracan evolution. PeerJ 7, e7387. <https://doi.org/10.7717/peerj.7387>

Feldmann, R.M., Villamil, T., Kauffman, E.G., 1999. Decapod and Stomatopod Crustaceans from Mass Mortality Lagerstatten: Turonian (Cretaceous) of Colombia. J. Paleontol. 73, 12.

Graham, J.R., Sevastopulo, G.D., 2021. The stratigraphy of latest Devonian and earliest Carboniferous rocks in Ireland. Palaeobiodiversity Palaeoenvironments. 101, 515-527. <https://doi.org/10.1007/s12549-020-00455-y>

Hancock, Z.B., Ogawa, H., Light, J.E., Wicksten, M.K., 2020. Origin and evolution of the Haustoriidae (Amphipoda): A eulogy for the Haustoriidira. BioRxiv. <https://doi.org/10.1101/2020.10.24.353664>

Hegna, T.A., Lazo-Wasem, E.A., Serrano-Sánchez, M. de L., Barragán, R., Vega, F.J., 2020a. A new fossil talitrid amphipod from the lower early Miocene Chiapas amber documented with microCT scanning. J. South Am. Earth Sci. 98, 102462. <https://doi.org/10.1016/j.jsames.2019.102462>

Hegna, T.A., Luque, J., Wolfe, J.M., 2020b. The fossil record of the Pancrustacea, in: Poore, G.C.B., Thiel, M. (Eds.), Evolution and Biogeography, Natural History of the Crustacea. Oxford University Press, Oxford.

Hilgen, F.J., Lourens, L.J., Van Dam, J.A., Beu, A.G., Boyes, A.F., Cooper, R.A., Krijgsman, W., Ogg, J.G., Piller, W.E., Wilson, D.S., 2012. The Neogene Period, in: The Geologic Time Scale. Elsevier, pp. 923–978.

Jarvis, D.E., 1990. New palynological data on the age of the Kiltorcan Flora of Co. Kilkenny, Ireland. J. Micropalaeontol. 9, 87-94. <https://doi.org/10.1144/jm.9.1.87>

Jarvis, D.E., 2000. Palaeoenvironment of the plant bearing horizons of the Devonian-Carboniferous Kiltorcan Formation, Kiltorcan Hill, Co. Kilkenny, Ireland. Geol. Soc. Spec. Publ. 180, 333-341. <https://doi.org/10.1144/GSL.SP.2000.180.01.16>

Lowry, J.K., Myers, A.A., 2017. A Phylogeny and Classification of the Amphipoda with the establishment of the new order Ingolfiellida (Crustacea: Peracarida). Zootaxa 4265, 1. https://doi.org/10.11646/zootaxa.4265.1.1

Lowry, J.K., Myers, A.A., 2013. A Phylogeny and Classification of the Senticaudata subord. nov. Crustacea: Amphipoda). Zootaxa 3610. <https://doi.org/10.11646/zootaxa.3610.1.1>

Lozano-Fernandez, J., Giacomelli, M., Fleming, J., Chen, A., Vinther, J., Thomsen, P.F., Glenner, H., Palero, F., Legg, D.A., Iliffe, T.M., Pisani, D., Olesen, J., 2019. Pancrustacean evolution illuminated by taxon-rich genomic-scale data sets with an expanded remipede sampling. Genome Biol. Evol. evz097. <https://doi.org/10.1093/gbe/evz097>

Luque, J., Feldmann, R.M., Vernygora, O., Schweitzer, C.E., Cameron, C.B., Kerr, K.A., Vega, F.J., Duque, A., Strange, M., Palmer, A.R., Jaramillo, C., 2019. Exceptional preservation of mid-Cretaceous marine arthropods and the evolution of novel forms via heterochrony. Sci. Adv. 5, eaav3875. <https://doi.org/10.1126/sciadv.aav3875>

Luque, J., Gerken, S., 2019. Exceptional preservation of comma shrimp from a mid-Cretaceous Lagerstätte of Colombia, and the origins of crown Cumacea. Proc. R. Soc. B Biol. Sci. 286, 20191863. <https://doi.org/10.1098/rspb.2019.1863>

Montagna, M., 2020. Comment on Phylogenetic analyses with four new Cretaceous bristletails reveal inter-relationships of Archaeognatha and Gondwana origin of Meinertellidae. Cladistics. 36, 227-231.

Montagna, M., Haug, J.T., Strada, L., Haug, C., Felber, M., Tintori, A., 2017. Central nervous system and muscular bundles preserved in a 240 million year old giant bristletail (Archaeognatha: Machilidae). Sci. Rep. 7, 46016. <https://doi.org/10.1038/srep46016>

Montagna, M., Tong, K.J., Magoga, G., Strada, L., Tintori, A., Ho, S.Y.W., Lo, N., 2019. Recalibration of the insect evolutionary time scale using Monte San Giorgio fossils suggests survival of key lineages through the End-Permian Extinction. Proc. R. Soc. B Biol. Sci. 286, 20191854.

Naumenko, S.A., Logacheva, M.D., Popova, N.V., Klepikova, A.V., Penin, A.A., Bazykin, G.A., Etingova, A.E., Mugue, N.S., Kondrashov, A.S., Yampolsky, L.Y., 2016. Transcriptome-based phylogeny of endemic Lake Baikal amphipod species flock: fast speciation accompanied by frequent episodes of positive selection. Mol. Ecol. <https://doi.org/10.1111/mec.13927>

Oakley, T.H., Wolfe, J.M., Lindgren, A.R., Zaharoff, A.K., 2013. Phylotranscriptomics to Bring the Understudied into the Fold: Monophyletic Ostracoda, Fossil Placement, and Pancrustacean Phylogeny. Mol. Biol. Evol. 30, 215–233. <https://doi.org/10.1093/molbev/mss216>

Robalino, J., Wilkins, B., Bracken-Grissom, H.D., Chan, T.-Y., O’Leary, M.A., 2016. The Origin of Large-Bodied Shrimp that Dominate Modern Global Aquaculture. PLOS ONE 11, e0158840. <https://doi.org/10.1371/journal.pone.0158840>

Robin, N., Gueriau, P., Luque, J., Daley, A.C., Vonk, R., 2021. The oldest peracarid crustacean reveals a Late Devonian freshwater colonization by isopod relatives. Biol. Lett. 17, 20210226. <https://doi.org/10.1098/rsbl.2021.0226>

Sánchez-Quiñónez, C.A., Tchegliakova, N., 2005. Foraminíferos planctónicos de la Formación San Rafael, Cretácico Superior, en los alrededores de Villa de Leiva, Boyacá, Colombia. Geol. Colomb. 30, 99–126.

Schwentner, M., Richter, S., Rogers, D.C., Giribet, G., 2018. Tetraconatan phylogeny with special focus on Malacostraca and Branchiopoda: highlighting the strength of taxon-specific matrices in phylogenomics. Proc. R. Soc. B Biol. Sci. 285, 20181524.

Serrano-Sánchez, M. de L., Hegna, T.A., Schaaf, P., Pérez, L., Centeno-García, E., Vega, F.J., 2015. The aquatic and semiaquatic biota in Miocene amber from the Campo LA Granja mine (Chiapas, Mexico): Paleoenvironmental implications. J. South Am. Earth Sci. 62, 243–256. <https://doi.org/10.1016/j.jsames.2015.06.007>

Solórzano Kraemer, M.M., 2007. Systematic, palaeoecology, and palaeobiogeography of the insect fauna from Mexican amber. Palaeontogr. Abt. A 282, 1–133.

Stockar, R., Baumgartner, P.O., Condon, D., 2012. Integrated Ladinian bio-chronostratigraphy and geochrononology of Monte San Giorgio (Southern Alps, Switzerland). Swiss J. Geosci. 105, 85-108. <https://doi.org/10.1007/s00015-012-0093-5>

Vega, F.J., Nyborg, T., Coutiño, M.A., Solé, J., Hernández-Monzón, O., 2009. Neogene Crustacea from Southeastern Mexico. Bull. Mizunami Foss. Mus. 35, 51–69.

Wolfe, J.M., Breinholt, J.W., Crandall, K.A., Lemmon, A.R., Moriarty Lemmon, E., Timm, L.E., Siddall, M.E., Bracken-Grissom, H.D., 2019. A phylogenomic framework, evolutionary timeline, and genomic resources for comparative studies of decapod crustaceans. Proc. R. Soc. B Biol. Sci. 286, 20190079. <https://doi.org/10.1101/466540>

Wolfe, J.M., Daley, A.C., Legg, D.A., Edgecombe, G.D., 2016. Fossil calibrations for the arthropod Tree of Life. Earth-Sci. Rev. 160, 43–110. <https://doi.org/10.1016/j.earscirev.2016.06.008>
